# Membrane potential correlates of network decorrelation and improved SNR by cholinergic activation in the somatosensory cortex

**DOI:** 10.1101/240853

**Authors:** Inbal Meir, Yonatan Katz, Ilan Lampl

**Author notes:** These authors equally contributed to this work (Inbal Meir, Yonatan Katz). Correspondence to: Ilan Lampl, Weizmann Institute of Science, Department of Neurobiology, 234 Herzl st., Rehovot 7610001, Israel.

## Abstract

The nucleus basalis (NB) projects cholinergic axons to the cortex where they play a major role in arousal, attention and learning. Cholinergic inputs shift cortical dynamics from synchronous to asynchronous and improves the signal to noise ratio (SNR) of sensory response. Yet, the underlying mechanisms of these changes remain unclear. Using simultaneous extracellular and whole cell patch recordings in layer 4 barrel cortex we show that activation of the cholinergic system has a differential effect on ongoing and sensory evoked activities. Cholinergic activation eliminated the large and correlated spontaneous synaptic fluctuations in membrane potential while sparing the synaptic response to whisker stimulation. This differential effect of cholinergic activation provides a unified explanation for the increased SNR of sensory response and for the reduction in both trial to trial variability and noise correlations as well as explaining the shift into desynchronized cortical state which are the hallmarks of arousal and attention.

## Introduction

The nucleus basalis (NB) plays a major role in arousal, attention, learning and plasticity (Chubykin et al., 2013; Froemke et al., 2012; Herrero et al., 2008; Kilgard and Merzenich, 1998; Xu et al., 2015). NB neurons send cholinergic, GABAergic and glutamatergic projections to the cortex (Gritti et al., 1997; Henny and Jones, 2008). During active states, such as locomotion or whisking cholinergic axons become active (Eggermann et al., 2014; Reimer et al., 2016). In-vivo NB activation or cholinergic modulation shift cortical dynamics from synchronous to asynchronous state, as seen in cortical EEG (Kalmbach et al., 2012; Metherate et al., 1992).

Acetylcholine was hypothesized to enhance the cortical sensory response by facilitation of thalamocortical inputs in L4, while suppressing the corticocortical inputs, thereby improving the signal to noise ratio (SNR) (Disney et al., 2007, 2012; Gil et al., 1997; Oldford and Castro-Alamancos, 2003). In contrast, other studies reported a cholinergic suppression of the sensory evoked spike response of L4 cells in the somatosensory and auditory cortex in vivo (Donoghue and Carroll, 1987; Sillito and Kemp, 1983), muscarinic suppression of both thalamocortical and intracortical synapses in L4 in the auditory cortex (A1) (Hsieh et al., 2000), and a persistent muscarinic hyperpolarization of L4 cells in S1, A1 and V1 in vitro (Eggermann and Feldmeyer, 2009).

Cholinergic modulation, NB activation, or attention, can further improve population sensitivity by neuronal spike decorrelation and by reducing noise correlations (Cohen and Maunsell, 2009a; Goard and Dan, 2009; Pinto et al., 2013a; Polack et al., 2013). The mechanisms underlying the increased SNR and the desynchronization of field potential and of neuronal firing, especially at the subthreshold level in vivo, remain unclear.

Using whole cell patch recordings and simultaneous local LFP in awake and anesthetized animals, we studied how global NB activation or local optogenetic activation of cholinergic fibers affect cortical layer 4 local network dynamics in the barrel cortex during ongoing and sensory-evoked activities. We have found that both types of stimulation had very similar effects on the cortical activity in L4. They suppressed the spontaneous activity, which kept the membrane potential close to the resting potential, rather than desynchronization of synaptic activity and depolarization of the membrane potential. The elimination of spontaneous activity served to increase the SNR of the sensory response and reduce both the spontaneous and noise-correlations in the local network.

## Results

### NB stimulation desynchronizes cortical LFP and increases sensory response SNR

To find how activation of the NB affects the summed synaptic cortical activity we recorded LFP in the barrel cortex at a depth of 400-450um, which corresponds to layer 4, in lightly anesthetized mice. We electrically stimulated the ipsilateral NB (see Methods) (Fig 1A – schematic illustration of the method). The location of the stimulating electrode was later verified histologically by a lesion performed at the end of the experiment. An example is shown in Figure 1B, illustrating a clear lesion at the corresponding location of the NB.

**Figure 1.**
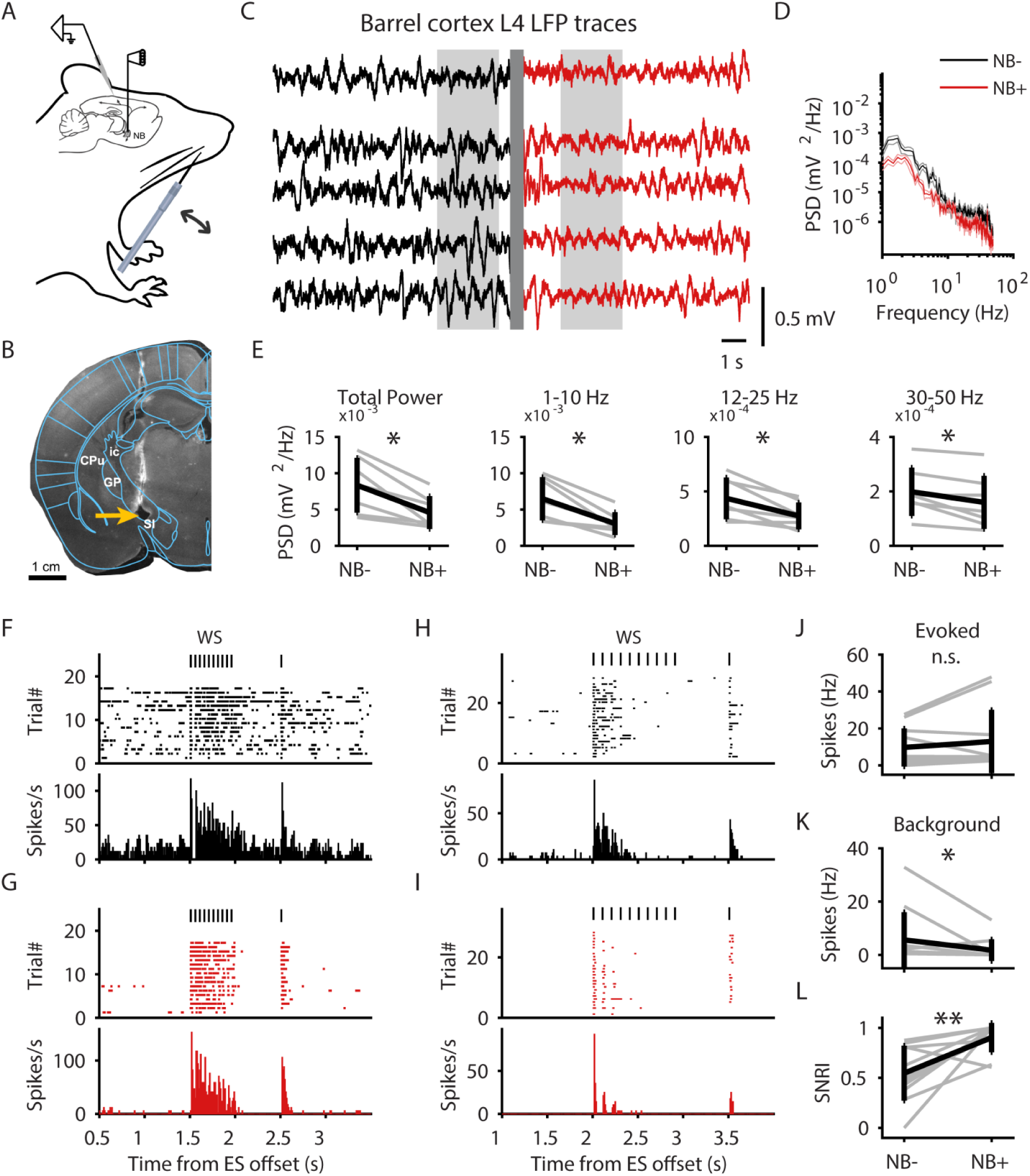
NB stimulation is shifting the cortex into an asynchronous state. (A) Schematic illustration of the method. Recordings in the barrel cortex were performed either with a single patch pipette or two pipettes for intracellular and nearby LFP recordings. (B) A lesion (orange arrow) marks the location of electrical stimulation electrode, corresponding to NB location. GP, Globus Pallidus, CPu, Caudate-Putamen, SI, Substantia Innominata (which contains the NB), ic, internal capsule. (C) Barrel cortex layer 4 LFP traces before (black; NB-) and after (red; NB+) electrical stimulation of the NB. Grey rectangles mark the part of the traces used to calculate the power spectral density (PSD) in (D-E). The trial averaged PSD is presented as mean±SEM. (E) from left to right: Total power is the sum of the Vm power spectrum, sum of power in frequencies 1-10Hz, sum of power in frequencies 12-25Hz, corresponding to beta band, sum of power in frequencies 30-50Hz, corresponding to lower gamma band. n=7 experiments. (F-I) The spontaneous and the sensory evoked response of two barrel cortex cells before (F,H) and after (G,I) NB stimulation. Upper panels: raster plots, Bottom panel: PSTHs. Time zero marks the offset of NB stimulation. (J) Rate of sensory evoked spikes during 500 ms of stimulus train. Background spike rate was subtracted. (K) Rate of background spikes during 500ms prior to sensory stimulation onset. (L) SNR index. In (E,J-L), grey lines mark single neurons, black lines mark population mean±SD. WS, whisker stimulation. * p<0.05, **p<0.01, Wilcoxon signed-rank test, n=11 cells.

Previous studies have shown that NB or cholinergic activation exerts both transient and prolonged modulation of cortical activity. The prolonged effect is most pronounced when inspecting the EEG or LFP activity which becomes desynchronized. This is manifested by elimination of slow waves and reduced spectral power in low frequencies (1-10 Hz) as well as an increase in gamma power (30-80 Hz). This effect could last several seconds (Kalmbach and Waters, 2014).

Consistent with the aforementioned studies, electrical stimulation at the corresponding depth of the NB caused an overall reduction in spectral power, especially in low frequencies (Figure 1C,D). Stimulating at other depths did not reduce the low-frequency power of cortical LFP, demonstrating the specificity of the effect to the location of the NB (Figure S1). For a population of LFP recordings from 7 experiments, we see a marked decrease in the total power (from 8.3·10^3^±3.7·10^3^ to 4.5·10^3^±2.2·10^3^ mV^2^/Hz, p=0.015, n=7, Figure 1E). This is a result of a general decrease in all 3 frequency bands tested: low frequency (1-10 Hz), beta (12-25 Hz) and gamma (30-50Hz) (p=0.015, p=0.015 and p=0.031, respectively, n=7 experiments, Figure 1E).

### NB stimulation increases the SNR of the sensory-evoked response

To study the effect of NB stimulation on the sensory-evoked response SNR we recorded in cell attached mode the spiking response of barrel cortical neurons to repetitive whisker deflection when it was either delivered 1.5 s after NB stimulation or without stimulating the NB.

Total of 11 cells were recorded in cell attached mode: 9 cells were recorded at depth of 350–600 μm, corresponding to layer 4 and 2 cells at 850-900 μm, corresponding to layer 5-6. The raster plots and PSTHs of two example cells are presented in Figure 1, showing a suppression of background spikes following NB stimulation while the magnitude of the evoked responses remained unchanged (Figure 1F-I).

To estimate the effect of NB stimulation on the response we calculated Response magnitude (see Methods). On average, there was no change in response modulation following NB stimulation (Figure 1J, from 9.72±10.3 to 12.95±17.11 spikes/s, p=0.17, Wilcoxon signed-rank test, n=11). Another way to quantify an NB-related modulation of the sensory-evoked response is by calculating a Modulation index (MI) (Vinck et al., 2015) (see Methods). The MI was not different than zero, indicating no change in the sensory evoked response following NB stimulation (Figure S2, MI = 0.034±0.31, p=0.76, n=11). However, the mean background activity (see Methods) was significantly reduced, reflecting the suppression of spontaneous firing (Figure 1K, from 5.64±10.42 to 1.78±4.08 spikes/s, p=0.02, n=11).

To quantify the Signal-to-Noise Ratio (SNR), we calculated an SNR index (Vinck et al., 2015) (see Methods). In line with previous studies, the SNR index showed a marked increase following NB stimulation (from 0.54±0.27 to 0.90±0.14, p=0.007, n=11, Figure 1L). Since there was no change in the Signal (the sensory response) the main contribution to the improved SNR is the reduced Noise (background firing).

Several mechanisms can account for the decrease in background spikes. Intrinsic changes can hyperpolarize the membrane or reduce the excitability of the cell by shunting; Neuromodulation can drive spiking of inhibitory cells or suppress firing of excitatory cells, or both. In order to better understand the underlying mechanisms, we made intracellular recordings.

### NB stimulation suppresses the spontaneous synaptic activity

We recorded the membrane potential of 17 barrel cortical cells from 13 animals using whole cell patch recording. All cells except for one were located at depths of 300-500um, corresponding to layer 4. An example from one cell is shown in Figure 2A-C. Membrane potential single traces demonstrate the effects of NB stimulation (Figure 2A). Following NB stimulation, the mean membrane potential became more hyperpolarized (Figure 2B) and the standard deviation decreased (Figure 2C). Across the population of cells, the mean membrane potential averaged along 1 second was hyperpolarized following NB stimulation from −59.59±7.89 mV to −61.84±7.87 mV (Figure 2D, Wilcoxon signed-rank test, n=17, p=0.0005). The trial-to-trial variability quantified by the membrane potential standard deviation decreased following NB stimulation from 4.89±1.99 mV to 3.55±2.05 mV (Figure 2E, Wilcoxon signed-rank test, n=17, p=0.0003).

**Figure 2.**
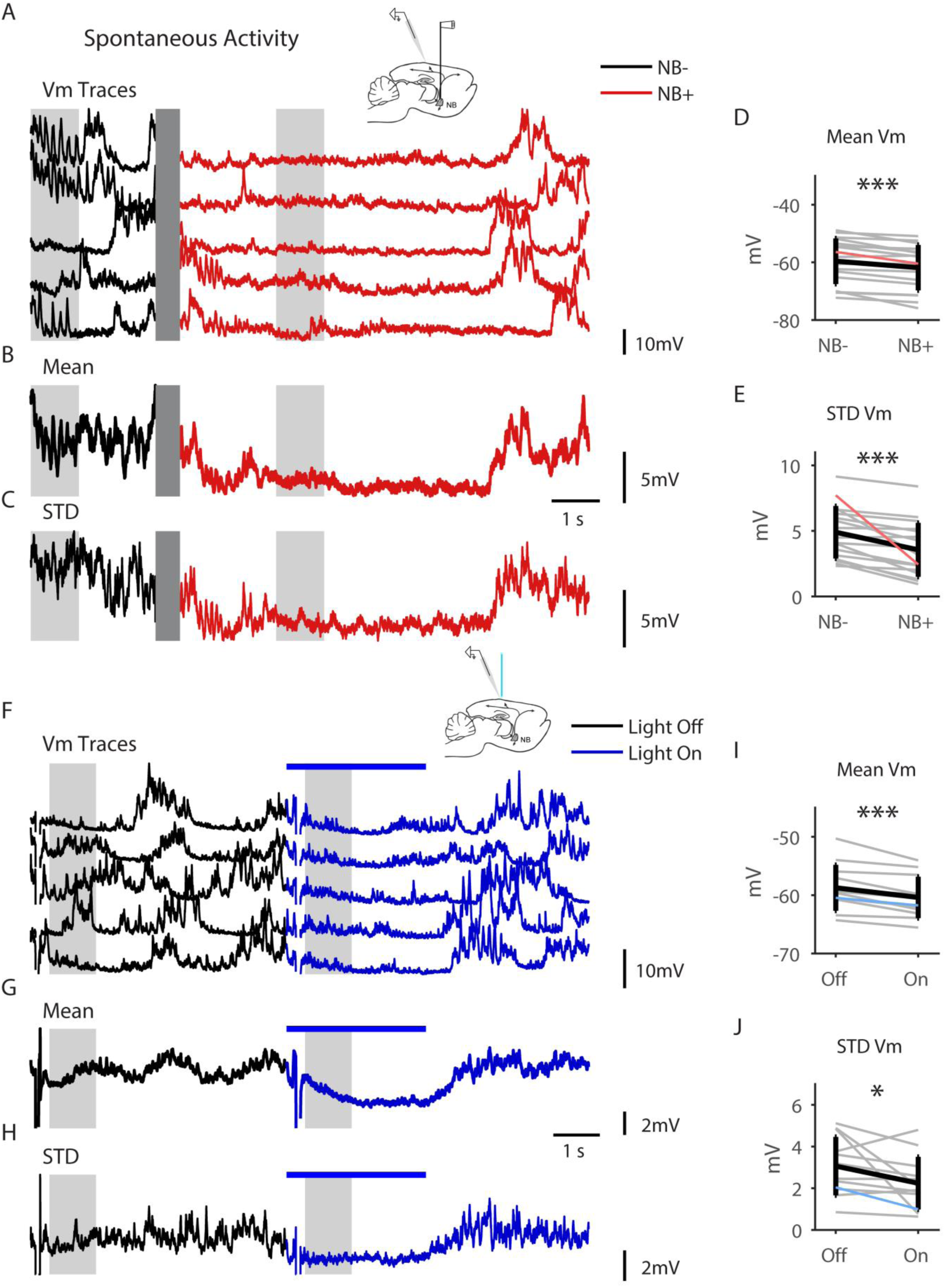
Cholinergic stimulation reduces the membrane potential mean and STD. (A-C) Spontaneous Vm traces (A), mean (B) and STD (C) of an example cell. (D) Population mean Vm. (E) Population Vm STD. (F-J) Same as (A-F) but before and during optogenetic stimulation. Grey lines mark single neurons (the example cell is in color), black lines mark population mean±SD. * p-value<0.05, *** p-value<0.001, (D,E) Wilcoxon signed-rank test, n=17 cells. (I, J) Wilcoxon signed-rank test, n=12 cells.

Electrical stimulation of the NB leads to global modulation due to the relatively wide axonal spread of NB cells. Moreover, the electrical stimulation non-selectively activates different NB cell populations, e.g. glutamatergic, cholinergic and GABAergic cells, along with a possible activation of fibers of passage or nearby cells. In order to stimulate only cholinergic inputs we used ChAT-ChR2 mice, which express ChR2 in cholinergic neurons (Figure S3A). First, we recorded extracellularly in the NB of ChAT-ChR2 mice using an optopatcher (Katz et al., 2013), which enables simultaneous intracellular or extracellular recording and light emission through the same pipette (see illustration in Figure S3B). We recorded in cell attached mode 3 single units that responded to light stimulation. An example from one cell is shown in Figure S3C-E. We stimulated the cell with different frequencies between 1-20 Hz and different pulse durations between 2-20 ms. The cell responded with a single spike locked to the light pulses, seen in the raw data trace (Figure S3C) and in raster- and PSTH-plots (Figure S3D,E). At frequencies above 5Hz the ability of the cells to follow the stimulus train decreased. It can be seen that at 15Hz the cell’s responses from the 3rd pulse onwards are less reliable. This is in accordance with previously reported characteristics of NB cholinergic cells (Kalmbach et al., 2012). Next, we recorded membrane potential using whole cell patch pipettes as described above and optogenetically activated the local cholinergic fibers innervating the cortex by 3 seconds continuous light stimulation emitted from a fiber optic placed <1mm above the cortical surface. We recorded the membrane potential of 12 barrel cortical cells in 5 animals. All cells were located at depths of 350-500um, corresponding to layer 4. An example of the effect of light stimulation on ongoing activity is shown in Figure 2F-H. Similar to the effects of NB activation with electrical stimulation, during light stimulation the mean membrane potential became more hyperpolarized (Figure 2G) and the standard deviation decreased (Figure 2H). In the population level, the mean membrane potential hyperpolarized significantly during light stimulation, from −58.73±3.91 mV to −60.38±3.54 mV (Figure 2I, p=0.005. Wilcoxon signed-rank test, n=12 cells). The trial-to-trial variability quantified by the membrane potential standard deviation decreased significantly during light stimulation from 3.06±1.4 mV to 2.24±1.26 mV. (Figure 2J, p=0.034. Wilcoxon signed-rank test, n=12). In contrast to the effect of NB electrical stimulation, here the changes in membrane potential dynamics started immediately after light onset and lasted at most 1 second after light offset. Some of the cells have shown a transient fast depolarization following light onset, matching nicotinic receptor activation.

As mentioned previously, the hyperpolarization of the mean membrane potential following NB stimulation may result from intrinsic mechanisms or due to synaptic network dynamics. Strong and prolonged inhibition would be required to suppress the large “bumps” of spontaneous activity. We expect such prominent shunting inhibition or shunting due to activation of intrinsic conductance to affect the input resistance of the cell. However, hyperpolarization of the mean membrane potential does not necessarily indicate that the recorded cells were directly inhibited. NB stimulation may reduce excitatory synaptic inputs, causing the distribution of membrane potential to shift towards the resting potential thus resulting with more negative mean membrane potential. Therefore, we examined the distribution of the membrane potential during spontaneous activity prior to and after NB stimulation. An example from one cell (same cell as in Figure 2A-C) is shown in Figure 3A. To estimate the true resting potential of the cell (i.e., the potential in absence of excitatory inputs) we calculated the lower 5^th^ percentile of Vm distribution. If NB stimulation was inhibiting the cell, we expect to find a shift in the 5^th^ percentile towards a more negative value. We found a mild decrease in the membrane potential 5^th^ percentile following NB stimulation (from −66±7.08 mV to −66.4±7.32 mV, p=0.039, n=17, Figure 3B). In 7 cells we injected a brief negative current pulse before and after NB stimulation and we did not see any change in the input resistance (from 41.44±30.44 MΩ to 44.84±21.16 MΩ, p=0.296, n=7, Figure 3C).

**Figure 3.**
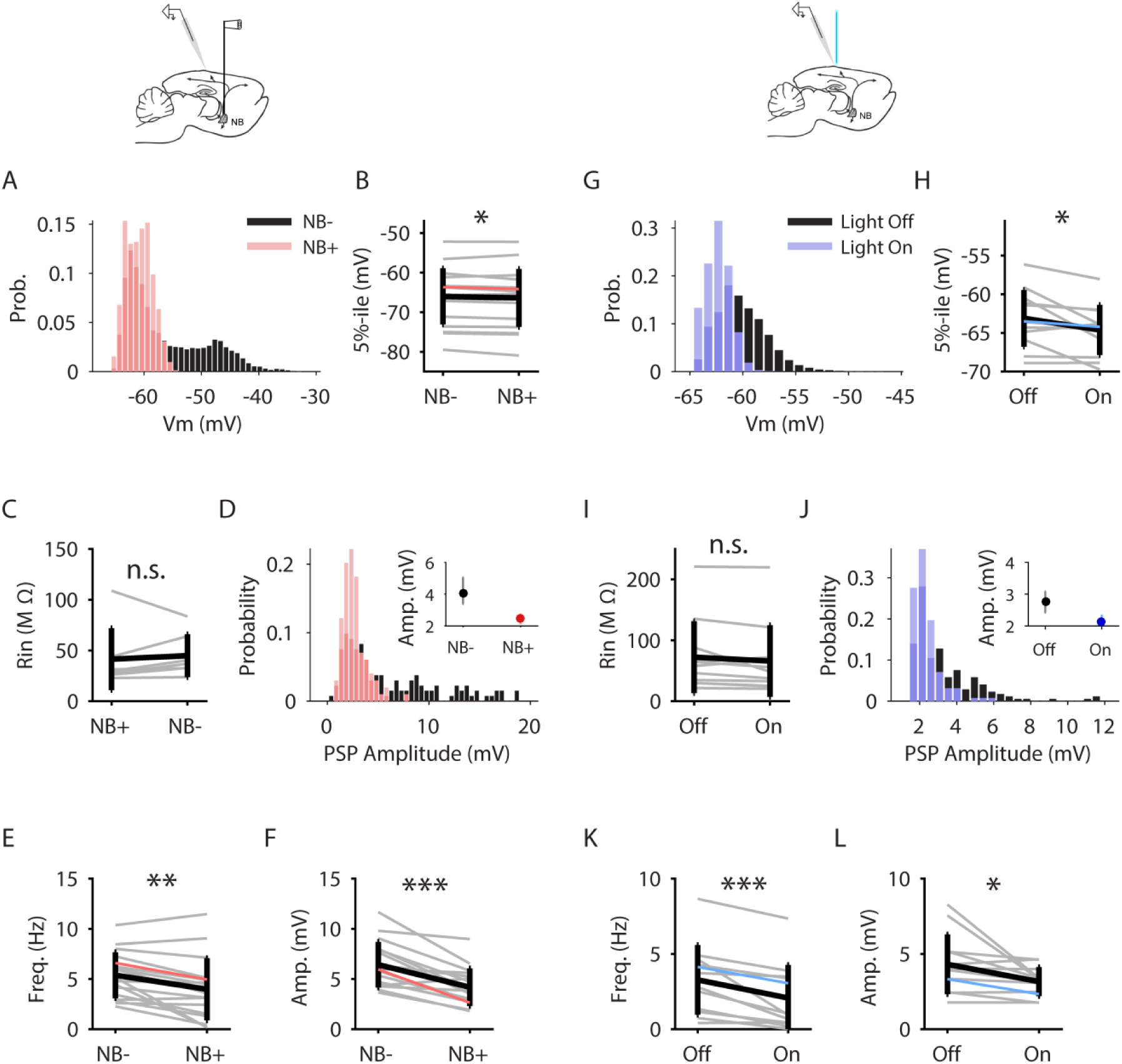
Cholinergic suppression of spontaneous synaptic activity. (A) Membrane potential histogram from one cell (same cell in Figure 2A-C). (B) Lower 5^th^ percentile of spontaneous Vm distribution. (C) Input resistance. (D) Histogram of PSPs amplitude from the same cell before (black) and after (red) NB stimulation. The inset shows the median amplitude for this cell with confidence intervals. (E) Frequency of spontaneous PSPs. (F) Amplitude of spontaneous PSPs. Grey lines mark single neurons (the example cell is in color), black lines mark population mean±SD. * p-value<0.05, ** p-value<0.01, *** p-value<0.001, Wilcoxon signed-rank test, n=17. (G-L.) Same as (A-F) but before and during optogenetic stimulation. Wilcoxon signed-rank test, n=12 cells.

Similar to NB stimulation, following light stimulation the lower 5^th^ percentile of the membrane potential distribution (an example of Vm histogram from one cell, same cell as in Figure 2F, is shown in Figure 3G) slightly decreased (from −63.12±3.67 mV to −155 64.62±3.24, p=0.016, Wilcoxon signed-rank test, n=12, Fig 3H), suggesting the possible involvement of inhibition or activation of intrinsic current. Yet, this process was not accompanied by any change in the input resistance (from 72.27±59.07 MΩ to 66.02±58.77 MΩ, p=0.12, n=11, Figure 3I). Together, the mild hyperpolarization of the membrane potential with no change in the input resistance indicate that some shunting inhibition is induced by NB activation but this inhibition is not strong enough to account for the elimination of the large spontaneous fluctuations in the membrane potential.

For a closer inspection of the spontaneous activity, we detected spontaneous post synaptic potentials (PSPs) (see Methods). Examples for detected spontaneous PSPs from four cells are shown in figure S4. The distribution of spontaneous PSPs from one cell before and after NB stimulation, demonstrates the decreased amplitude of spontaneous PSPs (Figure 3D). The inset shows the median PSP amplitude and confidence interval for this cell. Accordingly, the population analysis has revealed that following NB stimulation both the rate and the amplitude of spontaneous PSPs were diminished (from 5.37±2.29 Hz to 3.98±3.1 Hz, p=0.003, and 6.42±2.27 mV to 4.18±1.87 mV, p=0.0005, respectively, n=17, Figure 3E,F). We verified that the reduced rate of PSPs was not due to detection false negatives.

Similar to NB stimulation, a suppression of spontaneous activity during the light was indicated by a reduced rate and amplitude of spontaneous PSPs (from 3.27±2.31 Hz to 2.07±2.19 Hz, p=0.0009, and from 4.31±1.98 mV to 3.16±0.96 mV, p=0.016, respectively. Wilcoxon signed-rank test, n=12, Figure 3J-L). Taken together, our findings support a suppression of ongoing synaptic activity rather than an intrinsic mechanism or apparent synaptic inhibition.

### NB stimulation increases the SNR of subthreshold sensory responses

Next, we turned to characterize the effect of NB activation on the sensory evoked subthreshold responses in order to reveal the underlying mechanism for the improved response SNR shown in Figure 1. We delivered repeating trains of whisker stimuli (10 Hz or 20 Hz) with an additional identical test stimulus introduced 500 ms after the last stimulus in the train in order to examine if recovery from adaptation was affected by activation of cholinergic inputs. An example from a cell is shown in Figure 4A-D, demonstrating the effects of NB stimulation on sensory-evoked responses. Following NB stimulation, the mean membrane potential became more hyperpolarized, the sensory responses became more obvious on each stimulation (Figure 4C), and the trial-to-trial standard deviation decreased (Figure 4D). In the population level, the mean membrane potential during the sensory response hyperpolarized following NB stimulation from −54.35±8.13 mV to −55.71±8.48 mV (Figure 5A, Wilcoxon signed-rank test, n=17, p=0.0005). The trial-to-trial variability quantified by the membrane potential standard deviation during the sensory response decreased following NB stimulation from 4.22±1.57 mV to 3.76±1.89 mV (Figure 5B, Wilcoxon signed-rank test, n=17, p=0.031). Similar effects were seen during optogenetic activation of local cholinergic fibers, demonstrated in an example cell in Figure 4E-H. During light stimulation the mean membrane potential of the sensory response became more hyperpolarized (Figure 4G) and the standard deviation decreased (Figure 4H). In the population level, the mean membrane potential hyperpolarized significantly during light stimulation during the sensory train stimulus, from −56.5±4.69 mV to −58.45±4.24 mV, (Figure 5G, p=0.005. Wilcoxon signed-rank test, n=12 cells). The trial-to-trial variability, on the other hand, non-significantly decreased during light stimulation (from 2.36±0.62 mV to 2.25±0.9 mV during the sensory train Figure 5H, p=0.38. Wilcoxon signed-rank test, n=12). These changes seen in the membrane potential mean and standard deviation are similar to those seen during spontaneous activity, suggesting that NB stimulation or optogenetic activation of cholinergic fibers do not alter feedforward sensory inputs.

**Figure 4.**
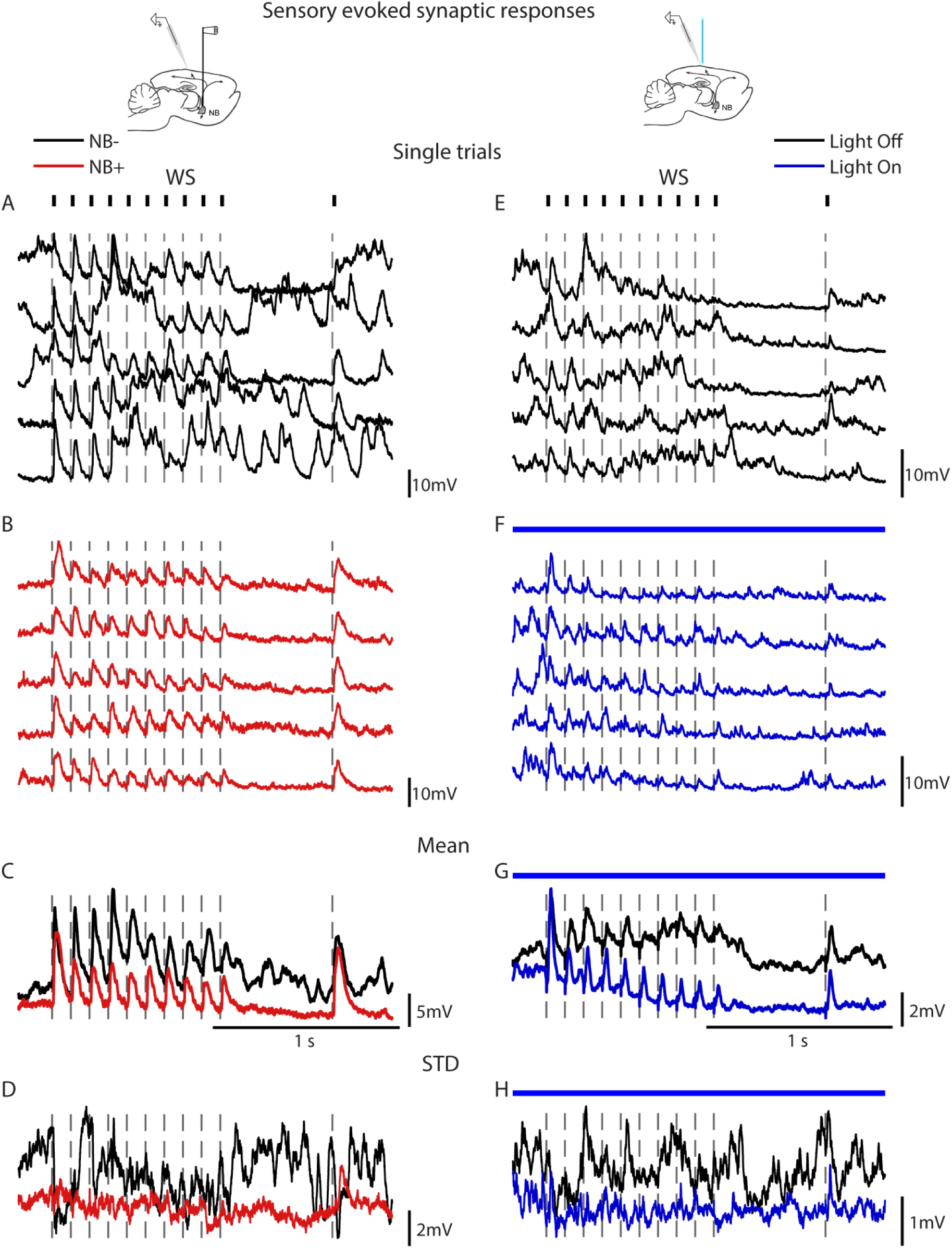
Cholinergic stimulation reduces the trial-to-trial variability. (A,B) Membrane potential traces of whisker evoked responses from an example cell before (A) and after (B) NB stimulation. (C) Mean trace. (D) Traces STD. (E-H) Same but in control and during optogenetic stimulation. WS, whisker stimulation. Dashed lines mark whisker deflections onset.

The effect of NB stimulation on the sensory response amplitude (baseline-to-peak) had complex dynamics. The response to the first stimulus was unchanged. In the rest of the sensory train stimulation, including the recovery response, there was a trend of reduced response, which was significant in the 3^rd^, 6^th^ and 8^th^ stimuli (Figure 5C, p=0.028, p=0.0057, p=0.0066, for stimuli 3,6, and 8, respectively. 0.1<p<0.7 for the rest of the stimuli in the train. two-way repeated measures ANOVA, n=17).

**Figure 5.**
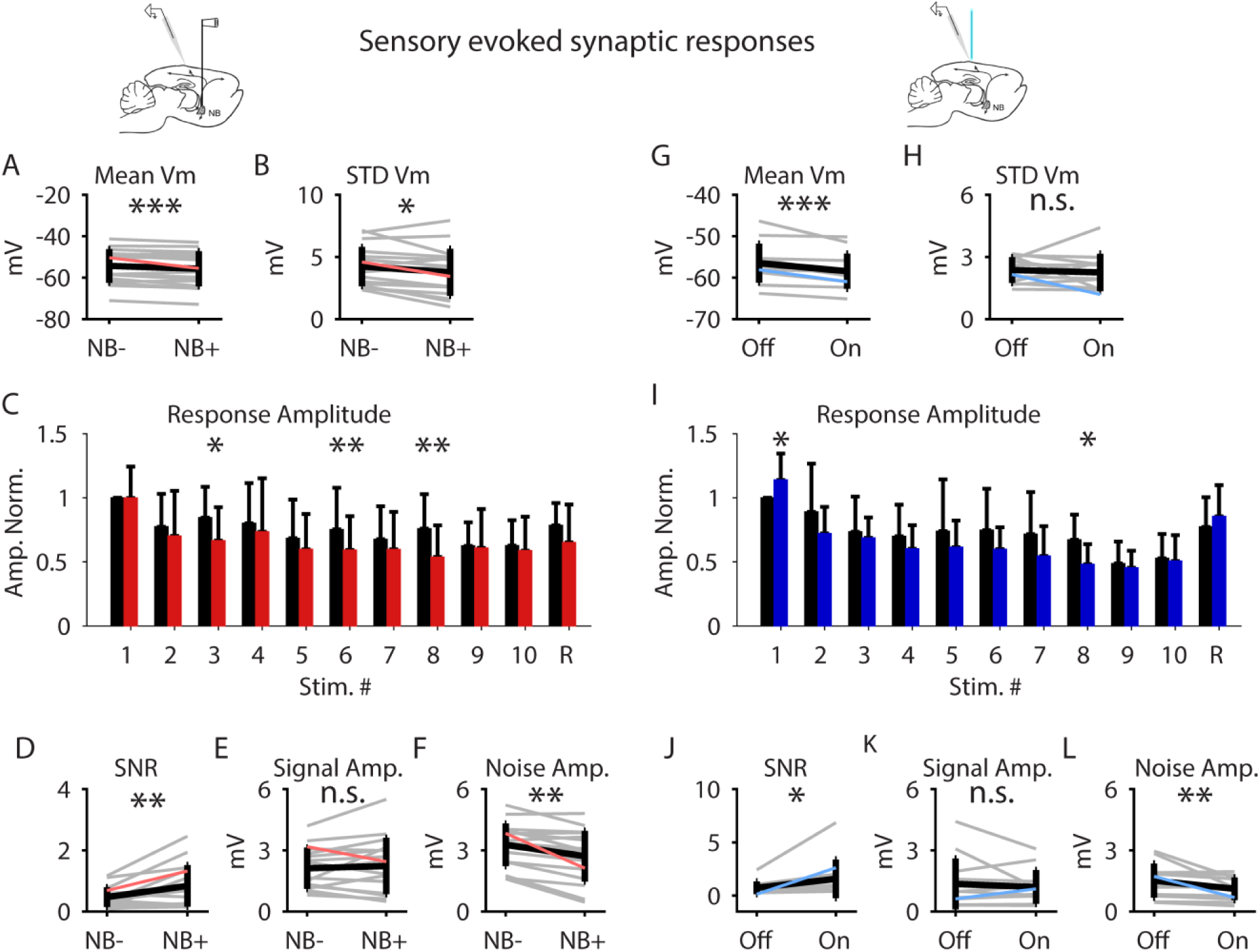
Cholinergic stimulation increases the SNR of subthreshold sensory responses. (A) Population mean membrane potential (Vm) whisker response. (B) Population Vm response STD. (C) Response amplitude along whisker stimulation train. Stimulus “R” marks recovery response. Lack of asterisk means no significant difference. (D) SNR. (E) Signal amplitude. (F) Noise amplitude. (G-L) Same as (A-F) but in control and during optogenetic stimulation. Grey lines mark single neurons (the example cell is in color), black lines mark population mean±SD. * p-value<0.05, ** p-value<0.01, *** p-value<0.001, n.s. non-significant, (C) Two-way repeated measures ANOVA, (A,B,D-F) Wilcoxon signed-rank test, n=17 cells. (I) Two-way repeated measures ANOVA, (G,H,J-L) Wilcoxon signed-rank test, n=12 cells.

The effect of light stimulation in ChAT-ChR2 mice on the subthreshold mean response amplitude was quite similar to the effect of NB electrical stimulation. There was a trend of reduced amplitude during the sensory stimulation train that reached significance only in the 8^th^ stimulus (from 4.02±3.96 mV to 2.71±2.78 mV, p=0.012, n=12, Figure 5I). In contrast to NB electrical stimulation effects, here the response to the first stimulus in the train was significantly increased (from 5.59±5.52 mV to 6.15±5.77 mV, p=0.032, n=12, Figure 5I), and a similar trend but not significant, was seen for the recovery response.

So far we have seen that activation of cholinergic inputs has differential effect on spontaneous and sensory evoked activities. Whereas spontaneous synaptic activity was reduced by activation of cholinergic inputs, the amplitude of the sensory response remained mainly unchanged, suggesting that the subthreshold SNR increased.

We computed the signal amplitude and noise amplitude of sensory responses (see Methods) and the squared ratio between them gave the SNR. Following NB stimulation the subthreshold SNR was increased by 42% (from 0.47±0.32 to 0.84±0.67; p=0.008, n=17, Figure 5D). In agreement with our extracellular results and the fact that the response amplitude remained mostly unchanged, this was due to decreased noise amplitude (from 3.27±1.04 mV to 2.71±1.24 mV, p=0.001, n=17, Figure 5F) while the signal amplitude remained unchanged (from 2.12±1.01 mV to 2.23±1.38 mV, p=0.72, n=17, Figure 5E).

Similar to NB activation, during light stimulation the noise amplitude decreased significantly (from 1.52±0.83 mV to 1.17±0.55 mV, p=0.005, Wilcoxon signed-rank test, n=12, Figure 5L) while the signal amplitude remained unchanged (from 1.35±1.25 mV to 1.2±0.82 mV, p=0.79, Wilcoxon signed-rank test, n=12, Figure 5K), resulting in an 115% increased subthreshold SNR (from 0.73±0.6 to 1.57±1.81 p=0.034, Wilcoxon signed-rank test, n=12, Figure 5J). In short, cholinergic modulation is increasing the sensory response SNR mainly by suppression of ongoing synaptic activity while only slightly affecting the sensory evoked responses.

### NB stimulation is decoupling the membrane potential and LFP signals and reducing the noise correlations

LFP amplitude is widely used as indication of population synchrony, where large amplitude LFP fluctuations are associated with high synchrony and a “flat” LFP is associated with a decorrelated population activity. In our recordings, we observed decreased LFP amplitude following NB stimulation along with decreased synaptic activity in individual cells. To find how activation of cholinergic input affects the correlation in the local network we made simultaneous recordings of membrane potential and LFP.

Simultaneous nearby Vm-LFP recordings (<500 μm) were recorded with electrical NB stimulation (n = 11) and with optogenetic stimulation (n = 9). An example of paired Vm-LFP recordings demonstrates high correlation between the signals during spontaneous and sensory evoked activities in the absence of cholinergic activation (Figure 6A, C, respectively, before NB stimulation, Figure 7A, C, before light stimulation). The two signals became much less correlated following activation of the NB (Figure 6B, D) or light stimulation (Figure 7B, D). To quantify the effect of NB or light stimulation on the correlation between the signals, we first estimated the correlations that result from either the NB or light stimuli as well as due to sensory stimulation. Towards this aim we generated sets of shuffled data traces where pairs of LFP and Vm traces were not recorded simultaneously and computed their cross-correlation (see Methods). In the case of responses to repeated sensory stimulus, the shuffled-correlations are in fact the “signal correlations” or “sensory-driven correlations”, while the actual data cross-correlation following subtraction of the shuffled correlations are the “noise correlations”. Such procedure was obtained also in the absence of sensory stimulation, when examining the effect of NB or light stimuli on the two signals and also when no stimulation at all was delivered. The shuffle-corrected Vm-LFP correlations from the same cell demonstrating a prominent decline in Vm-LFP correlation following NB or light stimulation during both spontaneous and sensory evoked responses (in Figure 6F, H and Figure 7F, H, the shuffled correlations are presented in Figure 6E,G and 7E, G).

**Figure 6.**
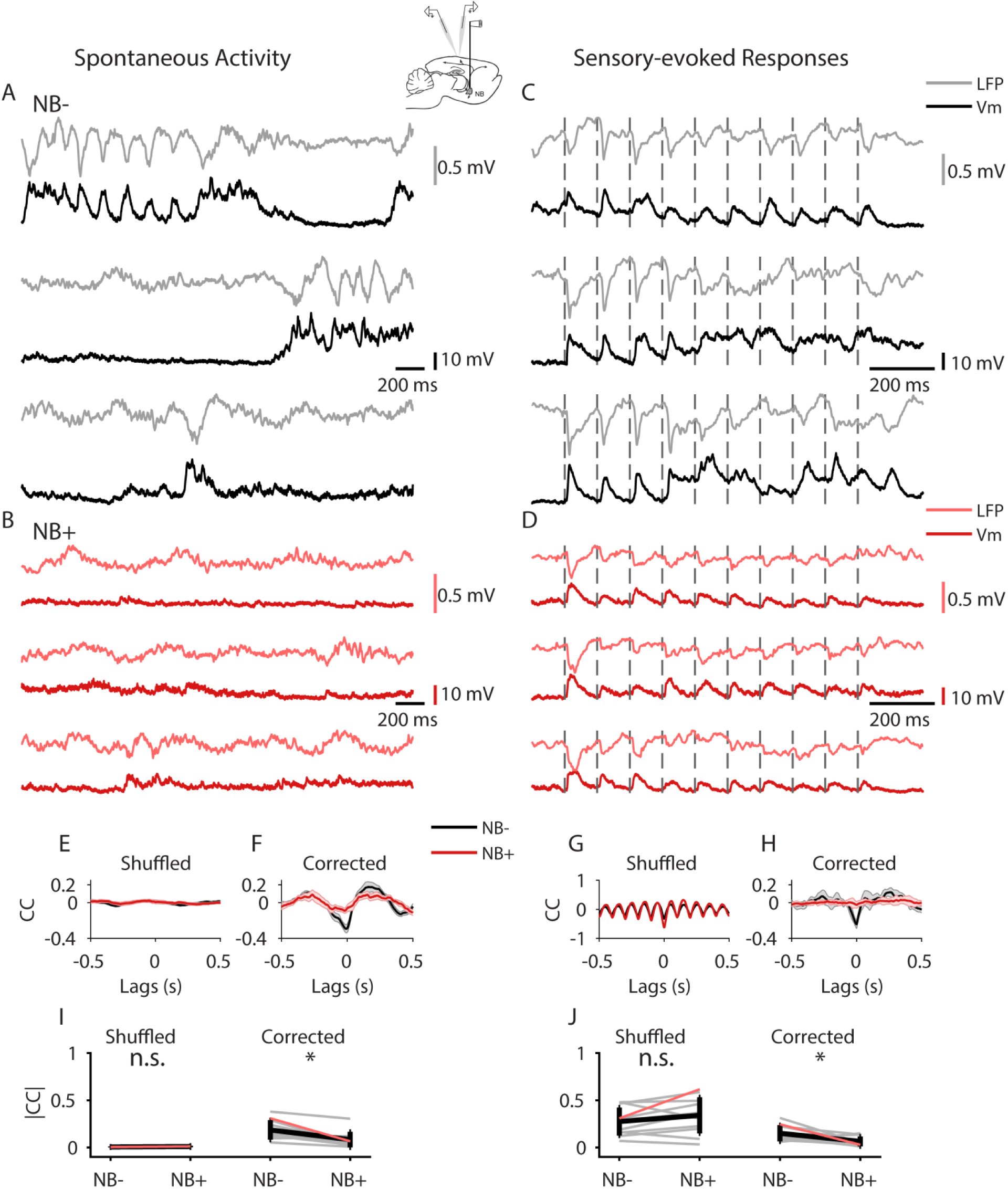
NB stimulation decorrelates the neuronal activity in the local cortical network. (A-H) Show an example from one cell. (A,B) Examples of 3 trace pairs of simultaneously recorded Vm and nearby LFP in control conditions (A) and following (B) NB stimulation, during spontaneous activity. (C,D) Same as (A,B) but for sensory-evoked responses. Dashed lines mark whisker deflections onset. (E) LFP-Vm shuffled traces cross-correlation. F. LFP-Vm cross-correlation in control conditions and following NB stimulation presented as mean±SEM. Shuffled correlations were subtracted. (G,H) Same as (E,F) but for sensory-evoked responses. (I) Population absolute-value Vm-LFP spontaneous zero-lag (Pearson) correlations. (J) Same as (I) but for sensory evoked responses. Grey lines mark single neurons (example cell is in color), black lines mark population average. * p-value<0.05, ** p-value<0.01, Two-way repeated measures ANOVA, n=11 cells.

**Figure 7.**
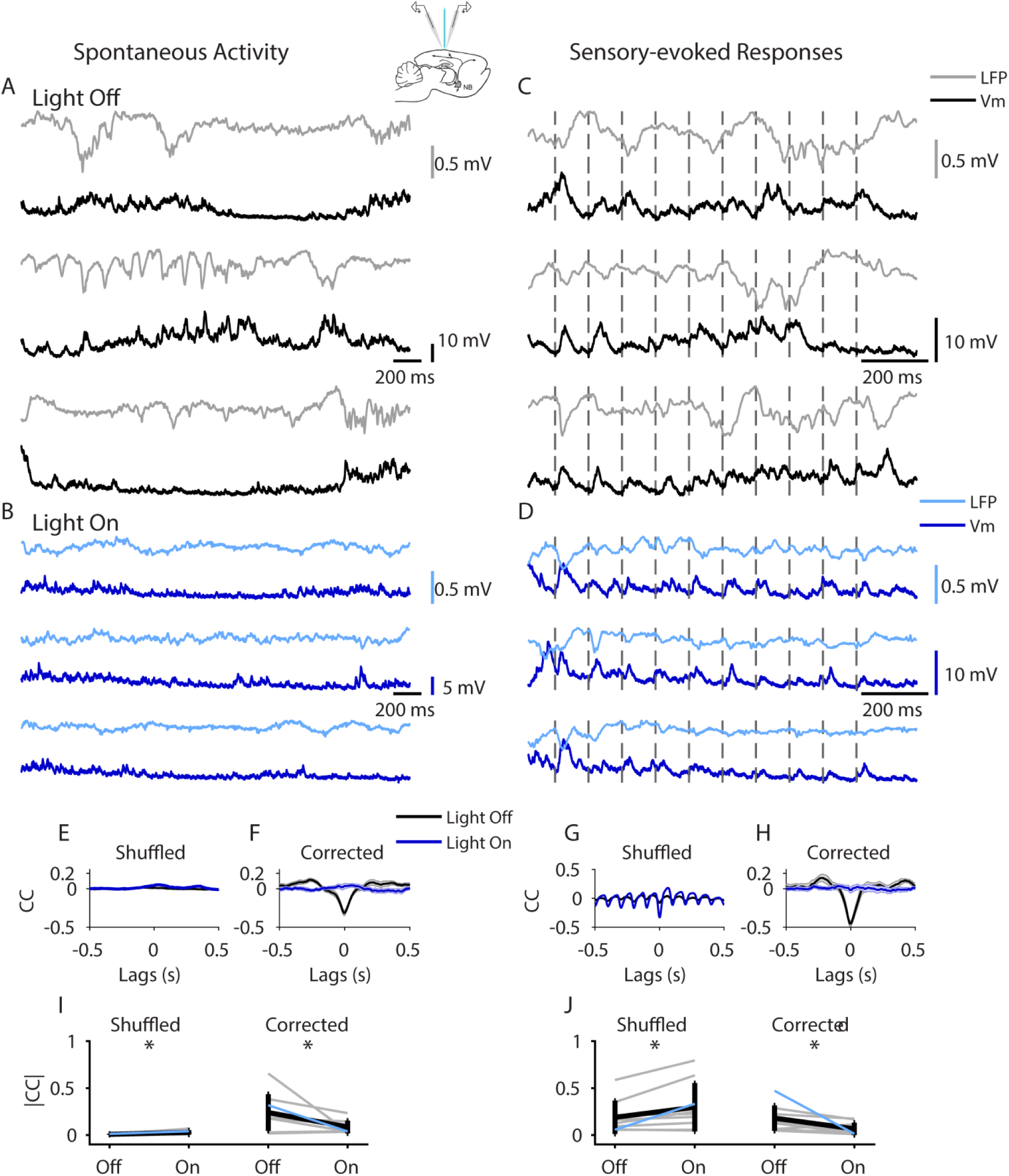
Cholinergic activation decorrelates the neuronal activity in the local cortical network. (A-H) Show an example from one cell. (A,B) Examples of 3 trace pairs of simultaneously recorded Vm and nearby LFP in control conditions (A) and during (B) light stimulation, during spontaneous activity. (C,D) same as (A,B) but for sensory-evoked responses. Dashed lines mark whisker deflections onset. (E) LFP-Vm shuffled traces cross-correlation. (F) LFP-Vm cross-correlation in control and during light stimulation presented as mean±SEM. Shuffled correlations were subtracted. (G,H) same as (E,F) but for sensory-evoked responses. (I) Population absolute-value Vm-LFP spontaneous zero-lag (Pearson) correlations. (J) Same as (I) but for sensory evoked responses. Grey lines mark single neurons (example cell is in color), black lines mark population average. * p-value<0.05, ** p-value<0.01, Two-way repeated measures ANOVA, n=9 cells.

In the population we found that NB stimulation or light stimulation caused marked reduction in the Vm-LFP correlations, almost to zero. In order to see this tendency, we took the absolute values of Vm-LFP zero time-lag correlations (Pearson correlations). During spontaneous activity the shuffled correlations (random correlations) were not significantly different following NB stimulation, indicating that NB stimulation did not introduced stimulus-locked activity in the network. However, the spontaneous shuffle-corrected correlations showed marked decline (Figure 6I, from 0.014±0.007 to 0.008±0.008, p=0.986, and from 0.192±0.128 to 0.077±0.086, p=0.038, for shuffled and shuffle-corrected correlations, respectively. Two-way repeated measures ANOVA with post-hoc Tukey comparisons, n=11). During the sensory evoked activity, shuffled correlations (sensory-driven correlations) did not change, while the shuffle-corrected correlations (noise correlations) significantly decreased following NB stimulation (Figure 6J, right panel, from 0.275±0.150 to 0.341±0.191, p=0.159, and from 0.150±0.087 to 0.063±0.051, p=0.013, for shuffled and shuffled-corrected correlations, respectively. Two-way repeated measures ANOVA with post-hoc Tukey comparisons, n=11).

The effects of light stimulation on the coupling between membrane potential and LFP were very similar. During light stimulation, the spontaneous shuffle-corrected correlations declined (Figure 7I, right panel, from 0.24±0.19 to 0.09±0.07, p=0.044) while the shuffled correlations did not change (Figure 7I, left panel, from 0.007±0.006 to 0.028±0.023, p=0.053, Two-way repeated measures ANOVA with post-hoc Tukey comparisons, n=9). During sensory stimulation, the noise correlations were reduced (right panel in Figure 7J), but in contrast to the case of NB stimulation – here the shuffled correlations during the sensory evoked activity (sensory-driven correlations) were increased (Figure 7J, from 0.18±0.17 to 0.3±0.25, p=0.031, and from 0.18±0.13 to 0.065±0.066, p=0.034, for shuffled and shuffled-corrected correlations, respectively. Two-way repeated measures ANOVA with post-hoc Tukey comparisons, n=9). The reduction in noise-correlations between Vm and LFP is consistent with previously reported decreases in neuronal spike correlations during attention, NB stimulation, or cholinergic modulation (Chen et al., 2015; Cohen and Maunsell, 2009b; Goard and Dan, 2009). The reduced spontaneous- and noise-correlations in the local neuronal network together with the suppression of spontaneous activity suggest that the ongoing activity prior to NB stimulation or light stimulation was correlated and thus introduced shared variability.

### Cholinergic suppression of spontaneous synaptic activity in awake animals

Anesthesia has profound effects on neuronal activity, especially in the cortex. To address the possibility that the ability of cholinergic inputs to modulate cortical activity is restricted to the state of anesthesia, we optogenetically activated local cholinergic fibers in the barrel cortex of animals that were awake and head-fixed. We recorded 7 L4 cells in 2 animals, while monitoring the animals’ locomotion and whisking (method illustrated in Figure 8A). In these experiments we only explored the effects of light on the spontaneous activity. An example from a cell is shown in Figure 8B-D, demonstrating the effects of light stimulation on the spontaneous activity. Similar to the anesthetized mice, the changes in membrane potential dynamics started immediately after light onset and lasted at most 1 second after light offset. Similar to the anesthetized preparation, we observed a suppression of spontaneous activity during light stimulation in the single trials (Figure 8B). The mean membrane potential became more hyperpolarized (Figure 8C) and the standard deviation decreased (Figure 8D). In the population level, the mean membrane potential hyperpolarized significantly during light stimulation, from −58.72±7.49 mV to −61.17±6.61 mV (Figure 8E, p=0.015. Wilcoxon signed-rank test, n=7 cells). The trial-to-trial variability quantified by the membrane potential standard deviation decreased significantly during light stimulation from 4.48±1.78 mV to 3.51±1.38 mV. (Figure 8F, p=0.015. Wilcoxon signed-rank test, n=7). To find if the reduction in mean membrane potential was only a result of suppressed ongoing activity or also by additional hyperpolarization we computed the distribution of membrane potential. A histogram of the membrane potential of the cell in 8B-D shows a shift of membrane potential distribution towards resting potential (Figure 8G). Similar to the results in the anesthetized mice, the 5^th^ percentile remained unchanged (−73.58±6.97 mV in control conditions and −72.47±6.2 mV during light, Figure 8I, p=0.218. Wilcoxon signed-rank test, n=7 cells).

**Figure 8.**
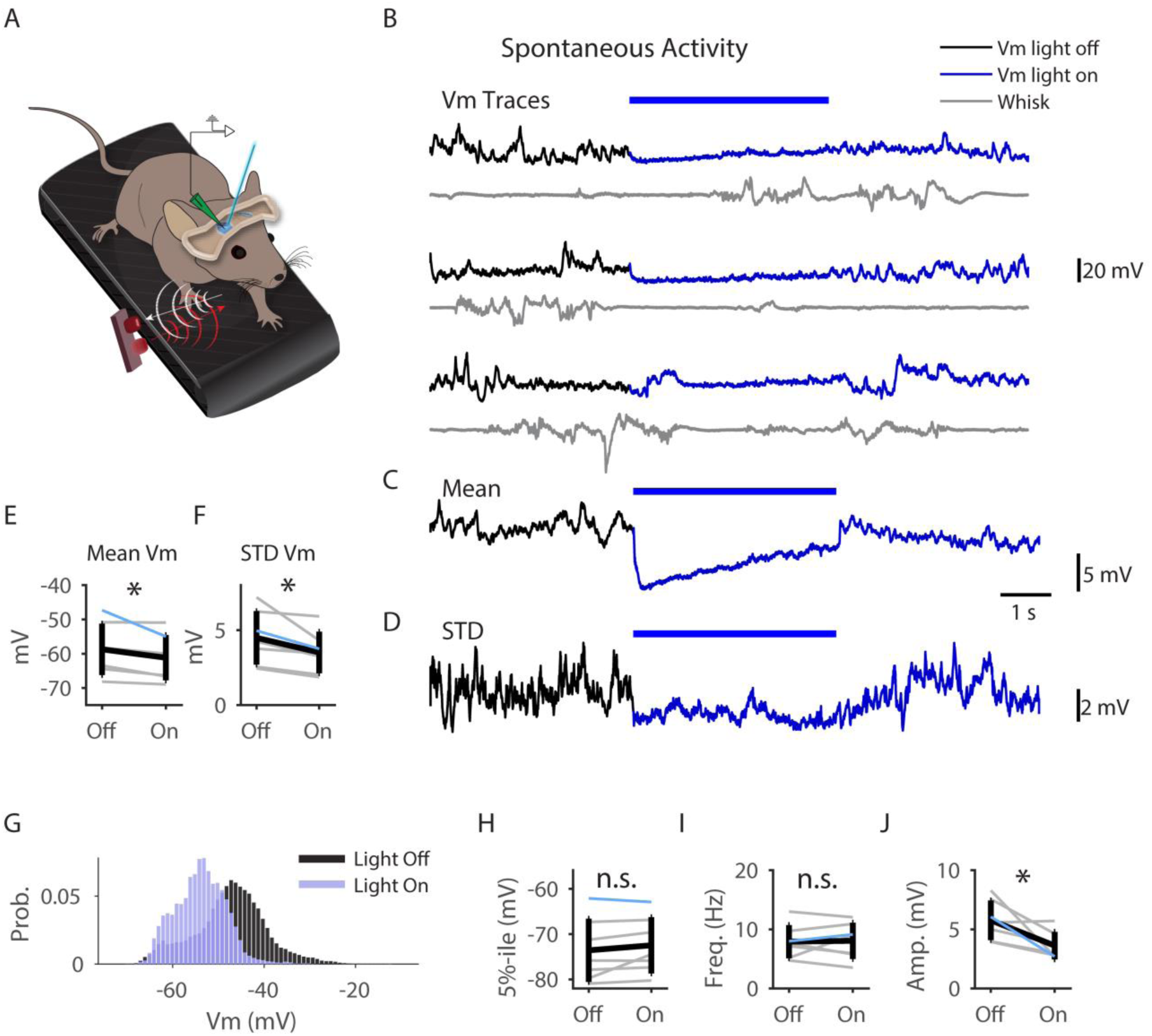
Cholinergic suppression of spontaneous synaptic activity in awake animals. (A) Illustration of recording and optogenetic stimulation method in awake head-fixed mice. (B-D) Spontaneous Vm (black) and whisking (grey) single traces recorded simultaneously (B), mean Vm (C) and STD Vm (D) of an example cell. (E) Population mean Vm. (F) Population Vm STD. (G) Membrane potential histogram from one cell (same cell in B-D). (H) Lower 5^th^ percentile of spontaneous Vm distribution. (I) Frequency of spontaneous PSPs. (J) Amplitude of spontaneous PSPs. Blue bar marks light stimulation. Grey lines mark single neurons (the example cell is in color), black lines mark population mean±SD. * p-value<0.05, ** p-value<0.01, *** p-value<0.001, n.s. non-significant, E,F,H,I,J Wilcoxon signed-rank test, n=7 cells.

To test the possibility that the shift in cortical state was caused by visual stimulation caused by the exposure of the animal to the light, we recorded from a WT animal. In 2 cells, we did not see any effect caused by the light (Figure S5). We also raised the possibility that seeing the light made the animal whisk and whisking shifted the cortical state. Comparing the average power of the whisking signal before and during the light stimulus showed that light did not induce whisking (Figure S6). Hence, our data suggests that the shift in cortical state induced by optogenetic stimulation of cholinergic projections in the barrel cortex of the awake brain resemble that seen during light anesthesia.

## Discussion

In this study, our aim was to better understand the synaptic and network mechanisms by which nucleus basalis neurons increase the signal to noise ratio (SNR) of sensory responses and decorrelate the cortical LFP. These effects are considered the main physiological correlates of attention and were so far studied, with very few exceptions, either intracellularly in vitro or extracellularly in vivo. However, the underlying membrane potential correlates of increased SNR of sensory response and network decorrelation were missing.

We have used cell attached and whole cell intracellular recordings as well as simultaneous nearby LFP recordings in both anesthetized and awake mice to investigate the firing and subthreshold modulation of somatosensory cortex layer 4 neurons by the nucleus basalis, either by electrically stimulating the NB or by optogenetically activating cholinergic axons which project to the somatosensory cortex.

We found that electrical stimulation of the NB or direct optogenetic activation of cholinergic cortical inputs increased the SNR of sensory responses by decreasing the ongoing synaptic activity while leaving the sensory response signal unaffected. These effects explain the reduction in the trial-to-trial variability of sensory responses and the reduced magnitude of ongoing activity. Likewise, reduced ongoing synaptic activity also explains the marked reduction in the correlation between the membrane potential and LFP during spontaneous-activity as well as the reduction in noise correlations of sensory evoked responses, while not affecting the sensory-driven signal correlations. Analysis of the cellular properties and membrane potential dynamics indicates that NB stimulation diminished rate and amplitude of spontaneous synaptic events, which led to a closer-to-rest membrane potential, with no change in input resistance. Importantly, we show that optogenetic activation of cholinergic fibers in awake mice results with very similar effects on ongoing activity as we found in anesthetized mice. Our findings therefore support the central role of the NB in the modulation of cortical circuits and strongly suggest that the major effect of cholinergic inputs is in suppression of ongoing cortical activity in the barrel cortex rather than altering feedforward tactile inputs.

That activation of the cholinergic inputs suppresses ongoing synaptic activity and leads to uncorrelated neuronal activity is in sharp contrast to the prolonged depolarization and increased firing associated with a “desynchronized” or “high-conductance” state (Destexhe et al., 2003; Renart et al., 2010). Thus, whereas reduced amplitude of LFP activity is usually regarded as one of the hallmarks of an active brain state, our combined Vm-LFP recordings indicate that reduced amplitude of LFP activity following cholinergic activation reflects in fact a marked drop in cortical activity.

### Cholinergic modulation of sensory response SNR

Cholinergic input has been long known to increase the SNR of the cortical sensory response. It is widely accepted that ACh enhances the sensory response by facilitating thalamocortical inputs in L4 while suppressing the intracortical inputs. Accordingly, this view is supported by in-vitro studies demonstrating that ACh can potentially modulate L4 neuronal activity by: (1) facilitation of sensory responses via presynaptic nicotinic receptors that enhance thalamocortical synaptic efficacy together with muscarinic presynaptic suppression of feedforward inhibition (2) muscarinic suppression of intracortical synapses, and further filtering by (3) depression due to post-synaptic muscarinic-induced prolonged intrinsic hyperpolarization (Eggermann and Feldmeyer, 2009; Gil et al., 1997; Hsieh et al., 2000; Kruglikov and Rudy, 2008).

Our study is the first to report the effect of cholinergic activation on subthreshold dynamics in response to a controlled sensory stimulus. Due to the non-homogenous spread of the cholinergic innervation and cholinergic receptors in the cortex, our use of NB electrical stimulation and especially local optogenetic activation of cholinergic axons mimics the physiological conditions of cholinergic modulation more closely than exogenous application of ACh or cholinergic agonists. Our findings are in line with a differential modulation of thalamocortical and intracortical pathways. We found very mild changes in sensory response amplitude with NB stimulation, and a modest enhancement of the subthreshold response amplitude during optogenetic cholinergic stimulation. In agreement with suppression of intracortical connections reported in vitro, we found a decreased rate and amplitude of spontaneous PSPs following NB activation. This suppression does not seem to be a result of direct muscarinic-mediated hyperpolarization as in (Eggermann and Feldmeyer, 2009) or synaptic inhibition, since there was no change in the input resistance and no clear net hyperpolarization (Figures 2 and 8). However, the exact mechanisms underlying this suppression of spontaneous activity are yet to be determined. Plausible mechanisms are (1) cholinergic depression of glutamate release from intracortical synapses (Gil et al., 1997) (2) dendritic inhibition via cholinergic activation of SOM interneurons (Muñoz et al., 2017) (3) suppression of cortical layer 5 (which we did not record in our study), which was shown to be the source of cortical spontaneous activity (Beltramo et al., 2013; Sakata and Harris, 2009) (4) cholinergic activation of L1 interneurons which inhibit pyramidal neurons through GABA_B_ (Brombas et al., 2014) or (5) long-lasting inhibition by cholinergic activation of astrocytes, which in turn excite inhibitory interneurons (Pabst et al., 2016). That ChR2-expressing cholinergic interneurons in layer 2/3 mediate these effects is unlikely. These neurons are scarce and they were shown to have negligible effects on neighboring cells (von Engelhardt et al., 2007). Our findings are at odds with the work of Metherate and Ashe (Metherate and Ashe, 1993) who have shown that NB electrical stimulation in vivo induced a persistent depolarization and increased excitability in the rat auditory cortex by closing a K^+^-mediated hyperpolarizing current. However, they recorded from more superficial layers than layer 4. Our results agree with the findings of Eggerman and colleagues (Eggermann et al., 2014), showing that optogenetic stimulation of cholinergic NB neurons suppress ongoing cortical activity in awake head-fixed mice. However, it is not clear why they have used only mice with inactivated thalamus. Last, in studies by (Pinto et al., 2013b) and (Minces et al., 2017), local optogenetic activation of cholinergic axons projecting to V1 has led to (1) increased spontaneous spikes across all layers in V1;(2) increased SNR due to increased signal amplitude with no change in the noise amplitude; (3) moderate decrease in noise correlations with no change in the signal correlations. The discrepancies between their work and our findings might be a result of the different cortical areas studied, but they might also be due to the use of different transgenic mouse lines. Pinto and colleagues used bacterial artificial chromosome (BAC) transgenic mice expressing channelrhodopsin-2 (*ChR2*) protein under the control of the choline acetyltransferase (*ChAT*) promoter (*ChAT–ChR2–EYFP mice*) which leads to overexpression of functional vesicular acetylcholine transporter gene (*VAChT)* and consequently increased cholinergic tone. It was demonstrated that these mice have severe cognitive deficits, including attention deficits and dysfunction in working memory and spatial memory, fundamentally differentiating them from wild-type mice (Kolisnyk et al., 2013).

The differential modulation of ongoing and sensory-evoked activities by NB activation observed in our study raises a difficulty. Recent studies have shown that the thalamic inputs account for only about a half of the sensory-evoked responses in layer 4 while the rest is provided by the cortical recurrent connections (Cohen-Kashi Malina et al., 2016; Lien and Scanziani, 2013), such that the sensory response is in fact composed of a thalamic and a cortical component. In that light, we would expect that suppression of the intracortical synapse due to cholinergic inputs would decrease the cortical component of the response and lead to an overall decreased response amplitude, but this was not the case. There might be a few explanations: 1) Facilitation of the thalamocortical inputs balances the muscarinic-induced suppression of the sensory response. 2) It might imply the existence of parallel intracortical circuits for ongoing activity and sensory-evoked responses, where the recurrent connections that give rise to the cortical component of the sensory response are spared while the connections that create the spontaneous fluctuations are suppressed by ACh.

Taken together, the finding that NB stimulation drastically reduces the noise amplitude but does not affect the signal amplitude, together with a negligible increase in subthreshold response amplitude but no change in response firing rate, highlight the cholinergic-induced noise reduction rather than signal enhancement.

### Different methods for activation of NB

The method of electrical stimulation for local activation has been used extensively in many brain areas, including the NB (Goard and Dan, 2009). This method has several disadvantages: 1) Stimulation artifacts may prevent us from analyzing the response during the stimulation. 2) Electrical stimulation activates NB cells regardless of cell type. 3) The effects on cortical activity might stem from activation of nearby cells or axons of passage, or by antidromic activation of axons that project to the NB and co-innervate the cortex or other structures that project to the cortex. 5) The modulation might be indirect, as electrical stimulation might activate NB cells which co-innervate or project to other brain areas which in turn innervate the barrel cortex. This can also happen with optogenetic activation, as optogenetic activation can generate antidromic spike in the ChR2-expressing axon that will spread to its collaterals. This is less likely as segregation of NB projections to different cortical areas has been demonstrated (Zaborszky et al., 2015). The difference in duration of the effect, which persists for longer time following electrical stimulation compared to optogenetic stimulation may be attributed to the global vs. local activation of the NB. Modulation of a larger cortical area might mean longer time for the ongoing activity to recover, e.g., less chances for stochastic events that will reinstate the recurrent activity in the network. Nevertheless, the fact that the two methods had such similar outcome points to a central role for ACh in the modulation of cortical circuits by the nucleus basalis.

### The effect of anesthesia

Anesthesia has profound effects on neuronal activity, especially in the cortex. Halothane potentiates GABA_A_ receptors and increases K^+^ leak currents, thereby hyperpolarizing the membrane potential (Rudolph and Antkowiak, 2004), in a manner that heavily depends on the dose. Our experiments were conducted both under light anesthesia and when animals were awake, demonstrating that the modulation of activity due to cholinergic activation was not related to anesthesia.

### Cortical Desynchronization

Active brain states such as arousal, attention and locomotion are physiologically characterized by desynchronized neuronal activity pattern and are also referred to as “high-conductance states” since it was suggested that the membrane conductance is high due to increased synaptic activity, both excitatory and inhibitory, which balance each other (Destexhe et al., 2003; Kumar et al., 2008; van Vreeswijk and Sompolinsky, 1996). High background activity in the high-conductance state depolarizes the membrane and increases the probability of a fluctuation to reach spike threshold and thus increase the spike rate. It was suggested that low spike correlations in this state are maintained through active decorrelation of synaptic currents by a fast tracking of excitation by inhibition (Renart et al., 2010). Our data demonstrate dissociation between the high-conductance state and the desynchronized state. In our experiments, reduced Vm-LFP correlations following NB or cholinergic stimulation appear together with reduced background synaptic activity, hyperpolarization of mean membrane potential, and subsequently suppressed spontaneous firing. Interestingly, recent studies in wakeful mice reported that during periods of high arousal without locomotion, the membrane potential dynamics resembled those we see following NB stimulation: hyperpolarized Vm, reduced Vm variance and suppression of low-frequency LFP fluctuations with no increase in high frequency fluctuations. Furthermore, these studies have shown that this state is optimal for cue detection (McGinley et al., 2015; Reimer et al., 2014; Vinck et al., 2015). An interesting question is regarding the dynamics of excitation and inhibition in this state, as it was shown that the rate of spontaneous activity can affect the ratio between excitation and inhibition (Taub et al., 2013).

### Thalamic involvement

The thalamus is known to play a role in cortical state (Poulet and Petersen, 2008; Steriade et al., 1993). NB does not innervate the sensory nuclei in the thalamus (e.g. ventral posterior medial nucleus (VPM) in the somatosensory system and lateral geniculate nucleus (LGN) in the visual system), but projections from NB to the reticular thalamus exist and might be responsible for some of the effects presented in this work (Levey et al., 1991; Pita-Almenar et al., 2014). However, this possibility was ruled out for V1 in a study that used NB electrical stimulation similar to our method, by extracellular recordings in LGN (Goard and Dan, 2009). Hence, it is unlikely that the effects of NB activation on cortical activity in our experiments were mediated via the thalamus.

### Trial-to-trial variability and noise correlations

The reliability of information encoding is affected by the trial to trial response variability of neuronal signals (Pillow et al., 2008; Zohary et al., 1994). Variability of sensory evoked cortical responses can be either private to a cell or shared among neurons. Private variability, e.g. due to internal processes in the cell such as the spike generation or external factors such as independent synaptic noise of unitary inputs (Deweese, 2004), can be averaged out if the population activity, rather than single neuron activity, is considered. Shared variability will lead to highly correlated trial-to-trial variability between neurons, i.e. noise correlations, which will not average out and therefore, might be detrimental for sensory coding, depending on the specific structure of correlations in the network (Averbeck et al., 2006).

There is an ongoing debate regarding the source of shared variability in cortical networks. A large body of research supports the idea of a cortical origin for the cortical noise correlations (Schölvinck et al., 2015; Timofeev et al., 2000), in contrast to cortical variability inherited from shared thalamic inputs (Bruno, 2006; Sadagopan and Ferster, 2012). A recent study has provided direct evidence that in contrast to the common assumption, the shared variability in the spontaneous activity and sensory responses in barrel cortex layer 4 is not a result of fluctuating correlated thalamic inputs but rather, the intracortical inputs are highly correlated while the thalamic inputs are weakly correlated (Cohen-Kashi Malina et al., 2016). Furthermore, noise correlations are modulated by the behavioral state of the animal (Cohen and Maunsell, 2009a) as well as by a sensory stimulus (Tan et al., 2014).

Our data revealed that activation of NB inputs of cortical cells significantly reduced both the membrane potential trial-to-trial variability and Vm-LFP noise correlations, suggesting that this variability was a shared variability introduced into the local network by correlated PSPs. These findings, and in particular our optogenetic experiments, further support a cortical origin of noise correlations. Moreover, they indicate that NB is capable of an active suppression of cortical noise correlations by acting directly on the cortex. To the best of our knowledge this is the first demonstration of cholinergic decorrelation of neuronal activity in-vivo at the synaptic level.

In summary, we have shown a cholinergic suppression of correlated cortico-cortical inputs that reduces the trial-to-trial variability of the membrane potential and reduces the correlations in the local population synaptic activity. This mechanism can account for the increased sensory response SNR and the reduced LFP amplitude commonly reported in cholinergic and attentional modulation, and thus enhances coding of sensory information.

## Materials and Methods

### Animals

All surgical and experimental procedures were performed in accordance with the regulations of The Weizmann Institute Animal Care and Use Committee. Recordings were made on young adult mice of either sex (9–16 weeks old) housed up to five in a cage with a 12/12h dark/light cycle. In experiments involving optogenetic cholinergic activation, we used ChAT-ChR2(Ai32) mice, which were the product of lines B6;129S6-Chat^tm1(cre)Lowl^/J (ChAT-Cre; Jax 006410) and 129S-Gt(ROSA)26Sor^tm32(CAG-COP4*H134R/EYFP)Hze^/J (Ai32, Jax 012569) (Madisen et al., 2012). ChAT-Cre+ mice were crossed with ChR2^+/+^ mice to yield Cre^+^ ChR2^+/−^ offspring. These offspring were crossed to generate Cre^+^ChR2^+/+^ mice, which were used for experiments. In NB-stimulation experiments we used WT C57B animals.

### Anesthetized animal preparation

WT or transgenic mice (15-30gr) were initially anesthetized intraperitoneally with ketamine (100 mg/kg) and xylazin (5 mg/kg). After the initial induction of anesthesia, animals were tracheotomized following the application of a local anesthesia, lidocaine (1%), injected subcutaneously and then a small metal tube was inserted into the trachea. Anesthesia was maintained by means of halothane (0.5-1%) and body temperature was kept at 37°C using a heating blanket. Craniotomy was performed over the somatosensory cortex (“barrel cortex”, centered 1.3 mm posterior and 3.3 mm lateral from bregma). In NB activation experiments, another craniotomy was made over the NB (1 mm posterior and 1.7 mm lateral from bregma).

### Awake animal preparation

Animals underwent the implantation of a head-bar to allow awake head-fixed recordings as follows: following initial anesthesia in an induction chamber containing a mix of isoflurane and oxygen-enriched air, the animals were mounted in a stereotaxic device, and kept deeply anaesthetized with isoflurane 1-1.5 %, monitored by checking for lack of reflexes and breathing rate. The area of incision was treated with lidocaine and cleaned with iodine and 70% ethanol. The skullcap was exposed and cleaned. The skull above the barrel cortex (centered 1.3 mm posterior and 3.3 mm lateral from the bregma) was covered with silicon glue (Smooth-On, Inc., USA). A plastic head-bar was firmly affixed to the skull with dental acrylic (3M, Germany). Half an hour before the end of the surgery the animals were injected subcutaneously with Buprenorphine (0.1 mg/Kg) and Carpofen (5 mg/Kg).

Following a recovery period (4–7 days), the animals were anaesthetized in an induction chamber containing a mix of isoflurane and oxygen-enriched air. The animals were then mounted in a stereotaxic device and kept deeply anaesthetized. The silicon glue covering the skull over the barrel cortex was removed and a craniotomy was performed exposing the barrel cortex and leaving the dura intact. The brain was then covered in an agar layer (2% w/v) held in place with silicon glue and the animal was returned to the cage for a recovery period of 2 h. The animal was then returned to the set and head-fixed for the electrophysiological recordings.

### Electrophysiology

#### Whole cell patch recordings

Borosilicate micropipettes were pulled to produce electrodes with a resistance of 4–10 MΩ when filled with an intracellular solution containing the following (in mM): K-gluconate, 136; KCl, 10; NaCl, 5; HEPES, 10; MgATP, 1; NaGTP, 0.3; phosphocreatine, 10; mOsm, 310. In some of the recordings, QX-314 (2 mM) was added to prevent spikes. Recording depth was ranged between 300 and 500 μm below pia. Intracellular signals were acquired using MultiClamp 700B amplifier (Molecular Devices) and low-pass filtered at 3 kHz before being digitized at 10 kHz.

#### LFP recordings

For LFP recordings, patch micropipettes were filled with artificial cerebrospinal fluid containing (in mM): 124 NaCl, 26 NaHCO3, 10 glucose, 3 KCl, 1.24 KH2PO4, 1.3 MgSO4 and 2.4 CaCl2. The pipette was inserted to a depth of 400-450 μm from pia. The signal was band-passed at 1-160Hz before being digitized at 10KHz. LFP was referenced to a chlorinated silver wire that was inserted under the skin of the neck. In simultaneous recordings of LFP and membrane potential, LFP pipette tip was placed at horizontal distance <500 μm from the recorded cell.

#### Extracellular recordings

Extracellular recordings were performed using patch micropipettes filled with patch solution. Recordings in cell-attached mode were made in the barrel cortex with regular pipette holder, and in the nucleus basalis using an optopatcher (Katz et al., 2013) (A-M systems, WA USA). The signals were amplified using MultiClamp 700B amplifier, band-passed at 0.3-3 kHz and digitized at 10 kHz.

#### Histology and immunohistochemistry

At the end of the experiment, the animals were over-dosed with anesthesia and transcardially perfused with PBS followed by cold 2.5% w/v paraformaldehyde (PFA). The brain was removed and postfixed in PFA for 24 hrs, followed by cryoprotection with a 30% (w/v) sucrose solution. The brain was then sectioned in 30 μm thick coronal slices using a sliding microtome (Leica SM 2010R; Nussloch, Eisfeld, Germany). The slices were mounted with Immu-Mount (Thermo Scientific Shandon). In experiments where the NB was electrically stimulated, the slices were examined and images of NB lesion were acquired with a fluorescent stereo microscope (Leica M165 FC). To confirm specific expression of ChR2 in cholinergic neurons we performed ChAT immunohistochemistry. Brain slices were thoroughly washed with PBS and incubated for 1 h with blocking buffer (20% w/v normal horse serum, 0.2% v/v Triton X-100 in PBS). The buffer was washed out with PBS and the slices were then incubated overnight at 4°C with primary antibody (1:100 dilution, goat anti-ChAT IgG (catalog no. AB144P, Millipore), and mouse anti-GFP IgG (catalog no. MAB3580, Millipore)), washed and incubated for 1.5 h at room temperature with the secondary antibody (1:200 dilution, biotinilated Donkey Anti-Goat IgG (catalog no. 705-065-147 JAC, Jackson Laboratories), or Cy2 conjugated anti-mouse (catalog number 715-225-151 JAC, Jackson Laboratories)). Images were acquired with a confocal Microscope (Zeiss, LSA 880; Carl Zeiss Microscopy GmbH, Jena, Germany) and processed in Adobe Photoshop 7.0.

#### Sensory stimulation

Whiskers were trimmed to a length of 10-15 mm. The principal whisker was inserted into a 21G needle and mechanically stimulated by a train of 10 deflections (20 ms duration) precisely controlled by a galvanometer servo control motor (6210H; Cambridge Technology Inc., USA), used with a matching servo driver and a controller (MicroMax 677xx; Cambridge Technology Inc., USA), at 10 or 20 Hz. In the majority of experiments we introduced an additional “recovery” stimulus – a single whisker deflection that was delivered 500 ms past train offset. The displacement was measured off-line using an optical displacement measuring system (optoNCDT 1605, Micro-Epsilon), indicating that ringing was negligible. A fast-rising voltage command was used to evoke a fast whisker deflection with a constant rise time of ~1 ms followed by a 20 ms ramp-down signal. The stimulation velocity and the corresponding deflection amplitude (~50 mm s^−1^, 145 μm amplitude) were adjusted to evoke clear subthreshold responses in the cortical cells.

In experiments with awake head-fixed mice, whisker movements were monitored using a reflective sensor (HOA1405) mounted on articulated arm. The sensor was positioned half a centimeter from the animals head and enabled the detection of whisking epochs and their phase.

#### Electrical and Optogenetic stimulation

For NB-stimulation, we stereotaxically placed a concentric needle electrode (30G; Alpine Biomed ApS) in the nucleus basalis in the area that modulates the somatosensory cortex (1 mm posterior and 1.7 mm lateral and 4.5-5mm ventral from bregma). A train of 50 current pulses of 100-200 μA at 100 Hz, pulse duration 0.5 ms, was injected by an isolation unit (ISO-Flex; A.M.P.I. Instruments). The depth positioning of the bipolar electrode was adjusted at the beginning of each experiment based on the cortical LFP responses to the electrical stimulation. Once the stimulating electrode position was set, it remained the same throughout the experiment. The stimulating electrode location was marked for histology at the end of the experiment by a current-induced lesion (2 sec, 5mA × 5 times).

For light stimulation, we used an analog modulated blue DPSS laser (λ = 473 nm, Shanghai Dream Lasers Technology Co. Ltd., Shanghai, China) coupled to a multi-mode fiber (NA = 0.22, 200μm core). The intensity of the light was around 7 mW at the tip of the fiber. For local cortical stimulation, continuous light pulse 3 or 4 seconds long was given through a fiber that was placed less than 1mm above the surface of the brain. In recordings from the NB, an optopatcher (Katz et al., 2013) was inserted into the nucleus basalis in the area that modulates the somatosensory cortex (1 mm posterior and 1.7 mm lateral and 4.5-5 mm ventral from bregma). Light pulses of 10 ms at frequencies between 1-15Hz were delivered in the nucleus basalis.

Stimulation protocol was composed of 3 trials in each repetition: 1) NB/light stimulation alone. 2) Whisker stimulation alone. 3) NB/light+whisker stimulation. Due to the long-lasting effect of NB-stimulation, we recorded 10 s long trials, with inter-trial-intervals of 10 s, allowing the neuronal activity to return to baseline. Trials were pseudo-randomized. Each trial type was repeated at least eight times in each cell. In the majority of experiments, we noticed a “bump” of activity immediately following NB stimulation offset, manifested as a bump with negative polarity in the LFP, increased spontaneous spiking in cell attached recordings and large spontaneous Vm depolarizations in whole-cell recordings, which could last from 0.1 to 1 second. We were interested in the prolonged effects of NB stimulation and therefore: (1) We set the onset of sensory stimulation to 1.5-2 seconds following NB stimulation offset, after the membrane potential has stabilized. (2) The analysis of spontaneous activity described next was performed on time windows starting 2.6 s prior to NB stimulation onset and 2.1 s after NB stimulation offset. Due to stimulation artifacts we were unable to analyze the modulation of LFP during the 500 ms of NB electrical stimulation (masked by a gray bar in Figure 1C).

#### Data Analysis

The data were analyzed using custom software written in MATLAB (The MathWorks).

In cell attached recordings, raster plots were binned at 10 ms bins to produce PSTHs.

Response magnitude was calculated from the PSTHs as follows:

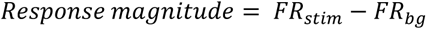

Where *FR_stim_* is the average firing rate during 500 ms of sensory train stimulus and *FR_bg_* is the background averaged firing rate during 500 ms immediately prior to the sensory stimulus. By subtracting the rate of background spikes from that during stimulation we estimated the net response to sensory stimulation, hence, the response magnitude which we refer to as the Signal.

Modulation index (MI) (Vinck et al., 2015) was calculated as follows:

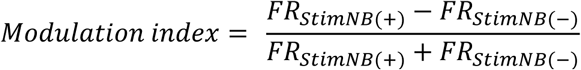

Where *FR_stimNB(−)_* is the firing rate during 500 ms train stimulation in absence of NB stimulation and *FR_stimNB(+)_* is the average firing rates during 500 ms following NB stimulation.

Mean background activity is *FR_bg_*, the background averaged firing rate during 500 ms immediately prior to the sensory stimulus.

To quantify the Signal-to-Noise Ratio (SNR), as evaluated from the response and background firing, we calculated an SNR index, which ranges between −1 and 1 (Vinck et al., 2015).

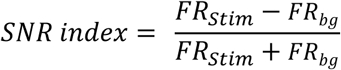

Where *FR_stim_* and *FR_bg_* are the same parameters as used when calculating *Response magnitude* (both for NB(+) and NB(−)).

In whole-cell recordings, some of the cells were spiking and the spikes were removed offline by spline interpolation based on 5 samples on each side of the action potential.

For calculation of membrane potential SNR, we used a definition of SNR of a waveform signal as the ratio of signal power and noise power. In the case where we are interested in the amplitude around the mean of the waveform and not relative to zero, such as our case, the SNR is equivalent to the ratio of signal variance and noise variance (Smith, 1997).

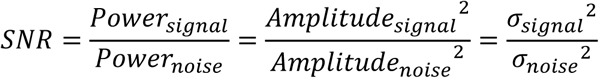

We took the signal to be the mean-over-trials of the sensory-evoked response and the noise as the residuals, i.e., the fluctuations around the mean. Therefore, we calculated the amplitude of the signal and the noise. The amplitude of a waveform is given by:

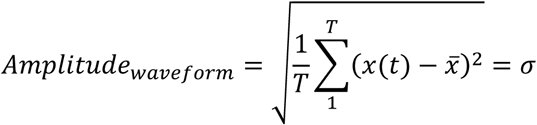

Where 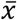 is the mean voltage over time, x(t) is the voltage at each time point t, and T is the total number of samples in the trace. For the noise amplitude, we calculated the amplitude of each residuals-trace and then averaged over traces. Finally, we squared both signal amplitude and noise amplitude. The ratio between them is the SNR.

In order to quantify the effect of NB stimulation or optogenetic stimulation on the membrane potential mean and trial-to-trial variability, we calculated the spontaneous and sensory-evoked Vm mean and standard deviation (STD). The mean or STD over trials was calculated and then averaged over time. The following time windows were used for the calculation: 1) Spontaneous Vm mean and STD were calculated from trials in which no sensory stimulus was presented, in time windows 1 s long starting 2.6 s prior to NB stimulation onset and 2.1 s after NB stimulation offset or 4.6 s prior to light onset and 0.4 s past light onset. 2) Sensory-evoked Vm mean and STD were calculated from 50 ms long time windows following each stimulus in the train, and then averaged over the stimuli in the train. The calculated values were compared between the two conditions – with and without NB stimulation or light stimulation.

Spontaneous and sensory-evoked post-synaptic potentials (PSPs) amplitude was measured as the difference between PSP peak and PSP onset values. We detected the events by using the derivative of the membrane potential to identify the onset- and peak-points. Events whose amplitude was greater than 1.5 mV were considered post synaptic potentials (PSPs) rather than noise (Okun and Lampl, 2008). This amplitude is somewhat above the ~1mV amplitude of noise fluctuations in the membrane potential recordings observed in raw data traces at the resting potential. The correct detection of events was verified manually to ensure optimal detection, with emphasis on minimizing false positives while slightly compromising false negative, i.e. missed events.

Histograms of membrane potential were made with 1 mV bins. Histograms of PSP amplitude were made with 0.5 mV bins. Confidence interval for the median PSP amplitude was computed using bootstrap method. We resampled the data and generated 5000 datasets. We calculated the median of each dataset and from these values took the 2.5 percentile and the 97.5 percentile to be the 95% confidence interval of the median.

For spectral analysis of LFP, we took the mean-subtracted traces from trials in which no sensory stimulation was introduced. We chose a 2.5 s long window starting 2.1 s past the offset of NB stimulation or 0.4 s past the onset of light stimulation, and a window of the same duration starting 2.6 s prior to NB stimulation onset or 4.6 s prior to light stimulation onset. The power spectral density (PSD) is the square of the fast Fourier transform (FFT), over frequency range between 1-49 Hz.

In the simultaneous Vm and LFP recordings, when power-line noise was obvious both signals were band-stop filtered at 49-51 Hz. For the cross-correlations of the spontaneous activity, we took two intervals of 2.5 s duration, starting 2.6 s prior to NB stimulation onset or 4.6 s prior to light stimulation onset, and 2.1 s after NB stimulation offset or 0.4 s past light stimulation onset. For sensory-evoked activity, as all cells in this analysis were sensory stimulated at 10 Hz, we took an interval of 1 s starting at the onset of the sensory stimulation train. Trials were mean-subtracted before computing the cross-correlation. For population analysis, we took the absolute-value of the zero-lag cross-correlation (Pearson correlation), averaged over trials. For each cell we generated 60 sets of shuffled data where pairs of LFP and Vm traces were not recorded simultaneously. These shuffled pairs underwent the same cross-correlation procedure as the actual data, averaged over the 60 iterations. These correlations were subtracted from the non-shuffled-correlations, yielding the shuffle-corrected correlations.

#### Statistical analysis

The statistical analyses for the populations was conducted using either Wilcoxon signed-rank test or repeated measures two-way ANOVA with Greenhause-Geisser correction if needed, followed by Tukey’s HSD post hoc tests. The significance level is at least p <0.05.

## Author contributions

Inbal Meir and Yonatan Katz: These authors equally contributed to this work.

I.M and I.L designed the research. I.M and Y.K conducted the experiments and analyzed the data, and I.M and I.L. wrote the manuscript.

## Acknowledgements

We would like to thank all the member of the Lampl lab for their helpful contribution to this work. This work was supported by grants DFG-SFB 1089, 01EW1606 - DeCipher EraNet Neuron, HFSP, Israel Science Foundation (ISF 1539/17) and Minerva, as well as by a research grant from the Marianne Manoville Beck Laboratory for Research in Neurobiology in Honor of her Parents Elisabeth and Miksa Manoville.

## References

1. Averbeck, B.B., Latham, P.E., and Pouget, A. (2006). Neural correlations, population coding and computation. Nat. Rev. Neurosci. 7, 358–366.

2. Beltramo, R., D’Urso, G., Dal Maschio, M., Farisello, P., Bovetti, S., Clovis, Y., Lassi, G., Tucci, V., De Pietri Tonelli, D., and Fellin, T. (2013). Layer-specific excitatory circuits differentially control recurrent network dynamics in the neocortex. Nat. Neurosci. 16, 227–234.

3. Brombas, A., Fletcher, L.N., and Williams, S.R. (2014). Activity-Dependent Modulation of Layer 1 Inhibitory Neocortical Circuits by Acetylcholine. J. Neurosci. 34, 1932–1941.

4. Bruno, R.M. (2006). Cortex Is Driven by Weak but Synchronously Active Thalamocortical Synapses. Science 312, 1622–1627.

5. Chen, N., Sugihara, H., and Sur, M. (2015). An acetylcholine-activated microcircuit drives temporal dynamics of cortical activity. Nat. Neurosci.

6. Chubykin, A.A., Roach, E.B., Bear, M.F., and Shuler, M.G.H. (2013). A Cholinergic Mechanism for Reward Timing within Primary Visual Cortex. Neuron 77, 723–735.

7. Cohen, M.R., and Maunsell, J.H.R. (2009a). Attention improves performance primarily by reducing interneuronal correlations. Nat. Neurosci. 12, 1594–1600.

8. Cohen, M.R., and Maunsell, J.H.R. (2009b). Attention improves performance primarily by reducing interneuronal correlations. Nat. Neurosci. 12, 1594–1600.

9. Cohen-Kashi Malina, K., Mohar, B., Rappaport, A.N., and Lampl, I. (2016). Local and thalamic origins of correlated ongoing and sensory-evoked cortical activities. Nat. Commun. 7, 12740.

10. Destexhe, A., Rudolph, M., and Paré, D. (2003). The high-conductance state of neocortical neurons in vivo. Nat. Rev. Neurosci. 4, 739–751.

11. Deweese, M.R. (2004). Shared and Private Variability in the Auditory Cortex. J. Neurophysiol. 92, 1840–1855.

12. Disney, A.A., Aoki, C., and Hawken, M.J. (2007). Gain Modulation by Nicotine in Macaque V1. Neuron 56, 701–713.

13. Disney, A.A., Aoki, C., and Hawken, M.J. (2012). Cholinergic suppression of visual responses in primate V1 is mediated by GABAergic inhibition. J. Neurophysiol.

14. Donoghue, J.P., and Carroll, K.L. (1987). Cholinergic modulation of sensory responses in rat primary somatic sensory cortex. Brain Res. 408, 367–371.

15. Eggermann, E., and Feldmeyer, D. (2009). Cholinergic filtering in the recurrent excitatory microcircuit of cortical layer 4. Proc. Natl. Acad. Sci. 106, 11753–11758.

16. Eggermann, E., Kremer, Y., Crochet, S., and Petersen, C.C.H. (2014). Cholinergic signals in mouse barrel cortex during active whisker sensing. Cell Rep. 9, 1654–1660.

17. von Engelhardt, J., Eliava, M., Meyer, A.H., Rozov, A., and Monyer, H. (2007). Functional characterization of intrinsic cholinergic interneurons in the cortex. J. Neurosci. Off. J. Soc. Neurosci. 27, 5633–5642.

18. Froemke, R.C., Carcea, I., Barker, A.J., Yuan, K., Seybold, B.A., Martins, A.R.O., Zaika, N., Bernstein, H., Wachs, M., Levis, P.A., et al. (2012). Long-term modification of cortical synapses improves sensory perception. Nat. Neurosci. 16, 79–88.

19. Gil, Z., Connors, B.W., and Amitai, Y. (1997). Differential Regulation of Neocortical Synapses by Neuromodulators and Activity. Neuron 19, 679–686.

20. Goard, M., and Dan, Y. (2009). Basal forebrain activation enhances cortical coding of natural scenes. Nat. Neurosci. 12, 1444–1449.

21. Gritti, I., Mainville, L., Mancia, M., and Jones, B.E. (1997). GABAergic and other noncholinergic basal forebrain neurons, together with cholinergic neurons, project to the mesocortex and isocortex in the rat. J. Comp. Neurol. 383, 163–177.

22. Henny, P., and Jones, B.E. (2008). Projections from basal forebrain to prefrontal cortex comprise cholinergic, GABAergic and glutamatergic inputs to pyramidal cells or interneurons. Eur. J. Neurosci. 27, 654–670.

23. Herrero, J.L., Roberts, M.J., Delicato, L.S., Gieselmann, M.A., Dayan, P., and Thiele, A. (2008). Acetylcholine contributes through muscarinic receptors to attentional modulation in V1. Nature 454, 1110–1114.

24. Hsieh, C.Y., Cruikshank, S.J., and Metherate, R. (2000). Differential modulation of auditory thalamocortical and intracortical synaptic transmission by cholinergic agonist. Brain Res. 880, 51–64.

25. Kalmbach, A., and Waters, J. (2014). Modulation of high- and low-frequency components of the cortical local field potential via nicotinic and muscarinic acetylcholine receptors in anesthetized mice. J. Neurophysiol. 111, 258–272.

26. Kalmbach, A., Hedrick, T., and Waters, J. (2012). Selective optogenetic stimulation of cholinergic axons in neocortex. J. Neurophysiol. 107, 2008–2019.

27. Katz, Y., Yizhar, O., Staiger, J., and Lampl, I. (2013). Optopatcher--an electrode holder for simultaneous intracellular patch-clamp recording and optical manipulation. J. Neurosci. Methods 214, 113–117.

28. Kilgard, M.P., and Merzenich, M.M. (1998). Cortical map reorganization enabled by nucleus basalis activity. Science 279, 1714–1718.

29. Kolisnyk, B., Guzman, M.S., Raulic, S., Fan, J., Magalhaes, A.C., Feng, G., Gros, R., Prado, V.F., and Prado, M.A.M. (2013). ChAT-ChR2-EYFP Mice Have Enhanced Motor Endurance But Show Deficits in Attention and Several Additional Cognitive Domains. J. Neurosci. 33, 10427–10438.

30. Kruglikov, I., and Rudy, B. (2008). Perisomatic GABA Release and Thalamocortical Integration onto Neocortical Excitatory Cells Are Regulated by Neuromodulators. Neuron 58, 911–924.

31. Kumar, A., Schrader, S., Aertsen, A., and Rotter, S. (2008). The high-conductance state of cortical networks. Neural Comput. 20, 1–43.

32. Levey, A., Kitt, C., Simonds, W., Price, D., and Brann, M. (1991). Identification and localization of muscarinic acetylcholine receptor proteins in brain with subtype-specific antibodies. J. Neurosci. 11, 3218–3226.

33. Lien, A.D., and Scanziani, M. (2013). Tuned thalamic excitation is amplified by visual cortical circuits. Nat. Neurosci. 16, 1315–1323.

34. Madisen, L., Mao, T., Koch, H., Zhuo, J., Berenyi, A., Fujisawa, S., Hsu, Y.-W.A., Garcia, A.J., 3rd, Gu, X., Zanella, S., et al. (2012). A toolbox of Cre-dependent optogenetic transgenic mice for light-induced activation and silencing. Nat. Neurosci. 15, 793–802.

35. McGinley, M.J., David, S.V., and McCormick, D.A. (2015). Cortical Membrane Potential Signature of Optimal States for Sensory Signal Detection. Neuron 87, 179–192.

36. Metherate, R., and Ashe, J.H. (1993). Ionic flux contributions to neocortical slow waves and nucleus basalis-mediated activation: whole-cell recordings in vivo. J. Neurosci. Off. J. Soc. Neurosci. 13, 5312–5323.

37. Metherate, R., Cox, C., and Ashe, J. (1992). Cellular bases of neocortical activation: modulation of neural oscillations by the nucleus basalis and endogenous acetylcholine. J. Neurosci. 12, 4701–4711.

38. Minces, V., Pinto, L., Dan, Y., and Chiba, A.A. (2017). Cholinergic shaping of neural correlations. Proc. Natl. Acad. Sci. U. S. A. 114, 5725–5730.

39. Muñoz, W., Tremblay, R., Levenstein, D., and Rudy, B. (2017). Layer-specific modulation of neocortical dendritic inhibition during active wakefulness. Science 355, 954–959.

40. Okun, M., and Lampl, I. (2008). Instantaneous correlation of excitation and inhibition during ongoing and sensory-evoked activities. Nat. Neurosci. 11, 535–537.

41. Oldford, E., and Castro-Alamancos, M.. (2003). Input-specific effects of acetylcholine on sensory and intracortical evoked responses in the “barrel cortex” in vivo. Neuroscience 117, 769–778.

42. Pabst, M., Braganza, O., Dannenberg, H., Hu, W., Pothmann, L., Rosen, J., Mody, I., van Loo, K., Deisseroth, K., Becker, A.J., et al. (2016). Astrocyte Intermediaries of Septal Cholinergic Modulation in the Hippocampus. Neuron 90, 853–865.

43. Pillow, J.W., Shlens, J., Paninski, L., Sher, A., Litke, A.M., Chichilnisky, E.J., and Simoncelli, E.P. (2008). Spatio-temporal correlations and visual signalling in a complete neuronal population. Nature 454, 995–999.

44. Pinto, L., Goard, M.J., Estandian, D., Xu, M., Kwan, A.C., Lee, S.-H., Harrison, T.C., Feng, G., and Dan, Y. (2013a). Fast modulation of visual perception by basal forebrain cholinergic neurons. Nat. Neurosci.

45. Pinto, L., Goard, M.J., Estandian, D., Xu, M., Kwan, A.C., Lee, S.-H., Harrison, T.C., Feng, G., and Dan, Y. (2013b). Fast modulation of visual perception by basal forebrain cholinergic neurons. Nat. Neurosci.

46. Pita-Almenar, J.D., Yu, D., Lu, H.-C., and Beierlein, M. (2014). Mechanisms underlying desynchronization of cholinergic-evoked thalamic network activity. J. Neurosci. Off. J. Soc. Neurosci. 34, 14463–14474.

47. Polack, P.-O., Friedman, J., and Golshani, P. (2013). Cellular mechanisms of brain state–dependent gain modulation in visual cortex. Nat. Neurosci.

48. Poulet, J.F.A., and Petersen, C.C.H. (2008). Internal brain state regulates membrane potential synchrony in barrel cortex of behaving mice. Nature 454, 881–885.

49. Reimer, J., Froudarakis, E., Cadwell, C.R., Yatsenko, D., Denfield, G.H., and Tolias, A.S. (2014). Pupil Fluctuations Track Fast Switching of Cortical States during Quiet Wakefulness. Neuron 84, 355–362.

50. Reimer, J., McGinley, M.J., Liu, Y., Rodenkirch, C., Wang, Q., McCormick, D.A., and Tolias, A.S. (2016). Pupil fluctuations track rapid changes in adrenergic and cholinergic activity in cortex. Nat. Commun. 7, 13289.

51. Renart, A., de la Rocha, J., Bartho, P., Hollender, L., Parga, N., Reyes, A., and Harris, K.D. (2010). The Asynchronous State in Cortical Circuits. Science 327, 587–590.

52. Rudolph, U., and Antkowiak, B. (2004). Molecular and neuronal substrates for general anaesthetics. Nat. Rev. Neurosci. 5, 709–720.

53. Sadagopan, S., and Ferster, D. (2012). Feedforward Origins of Response Variability Underlying Contrast Invariant Orientation Tuning in Cat Visual Cortex. Neuron 74, 911–923.

54. Sakata, S., and Harris, K.D. (2009). Laminar Structure of Spontaneous and Sensory-Evoked Population Activity in Auditory Cortex. Neuron 64, 404–418.

55. Schölvinck, M.L., Saleem, A.B., Benucci, A., Harris, K.D., and Carandini, M. (2015). Cortical state determines global variability and correlations in visual cortex. J. Neurosci. Off. J. Soc. Neurosci. 35, 170–178.

56. Sillito, A.M., and Kemp, J.A. (1983). Cholinergic modulation of the functional organization of the cat visual cortex. Brain Res. 289, 143–155.

57. Smith, S.W. (1997). The scientist and engineer’s guide to digital signal processing.

58. Steriade, M., McCormick, D.A., and Sejnowski, T.J. (1993). Thalamocortical oscillations in the sleeping and aroused brain. Science 262, 679–685.

59. Tan, A.Y.Y., Chen, Y., Scholl, B., Seidemann, E., and Priebe, N.J. (2014). Sensory stimulation shifts visual cortex from synchronous to asynchronous states. Nature 509, 226–229.

60. Taub, A.H., Katz, Y., and Lampl, I. (2013). Cortical balance of excitation and inhibition is regulated by the rate of synaptic activity. J. Neurosci. Off. J. Soc. Neurosci. 33, 14359–14368.

61. Timofeev, I., Grenier, F., Bazhenov, M., Sejnowski, T.J., and Steriade, M. (2000). Origin of slow cortical oscillations in deafferented cortical slabs. Cereb. Cortex N. Y. N 1991 10, 1185–1199.

62. Vinck, M., Batista-Brito, R., Knoblich, U., and Cardin, J.A. (2015). Arousal and Locomotion Make Distinct Contributions to Cortical Activity Patterns and Visual Encoding. Neuron 86, 740–754.

63. van Vreeswijk, C., and Sompolinsky, H. (1996). Chaos in neuronal networks with balanced excitatory and inhibitory activity. Science 274, 1724–1726.

64. Xu, M., Chung, S., Zhang, S., Zhong, P., Ma, C., Chang, W.-C., Weissbourd, B., Sakai, N., Luo, L., Nishino, S., et al. (2015). Basal forebrain circuit for sleep-wake control. Nat. Neurosci. 18, 1641–1647.

65. Zaborszky, L., Csordas, A., Mosca, K., Kim, J., Gielow, M.R., Vadasz, C., and Nadasdy, Z. (2015). Neurons in the basal forebrain project to the cortex in a complex topographic organization that reflects corticocortical connectivity patterns: an experimental study based on retrograde tracing and 3D reconstruction. Cereb. Cortex N. Y. N 1991 25, 118–137.

66. Zohary, E., Shadlen, M.N., and Newsome, W.T. (1994). Correlated neuronal discharge rate and its implications for psychophysical performance. Nature 370, 140–143.

